# Aberrant methylation and expression of TNXB promote chondrocyte apoptosis and extracullar matrix degradation in hemophilic arthropathy via AKT signaling

**DOI:** 10.1101/2023.12.11.570472

**Authors:** Jiali Chen, Zeng Qinghe, Xu Wang, Rui Xu, Weidong Wang, Yuliang Huang, Qi Sun, Wenhua Yuan, Pinger Wang, Di Chen, Peijian Tong, Hongting Jin

## Abstract

Recurrent joint bleeding in hemophilia patients frequently causes hemophilic arthropathy (HA). Drastic degradation of cartilage is a major characteristic of HA, but its pathological mechanisms has not yet been clarified. In HA cartilages, we found server matrix degradation and increased expression of DNA methyltransferase proteins. We thus performed genome-wide DNA methylation analysis on human HA (N = 5) and osteoarthritis (N = 5) articular cartilages, and identified 1228 differentially methylated regions (DMRs) associated with HA. Functional enrichment analyses revealed the association between DMR genes (DMGs) and extracellular matrix (ECM) organization. Among these DMGs, Tenascin XB (TNXB) expression was down-regulated in human and mouse HA cartilages. The loss of *Tnxb* in *F8^-/-^* mouse cartilage provided a disease-promoting role in HA by augmenting cartilage degeneration and subchondral bone loss. *Tnxb* knockdown also promoted chondrocyte apoptosis and inhibited phosphorylation of AKT. Importantly, AKT agonist showed chondroprotective effects following *Tnxb* knockdown. Together, our findings indicate that exposure of cartilage to blood leads to alterations in DNA methylation, which is functionally related to ECM homeostasis, and further demonstrate a critical role of TNXB in HA cartilage degeneration by activating AKT signaling. These mechanistic insights allow development of potentially new strategies for HA cartilage protection.

## Introduction

Hemophilia is a X-linked bleeding disorder due to the deficiency of coagulation factor VIII (hemophilia A) and IX (hemophilia B). As a one of the most frequent rare diseases, hemophilia is a persistent, challenging condition for patients, physicians and society ^1^. Joint bleeding (hemarthrosis), especially in knee joint, is the most common clinical manifestation in patients with severe hemophilia (i.e., plasma FVIII or FIX < 1 U/dl) ^2^. Recent evidence also indicates that hemarthrosis may occur spontaneously in patients with moderate (plasma factor levels of 1– 5 UI/dl) or mild disease (plasma factor levels of >5 UI/dl) ^3,4^. Repeated episodes of hemarthrosis eventually result in hemophilic arthropathy (HA), a debilitating and irreversible condition. HA often starts at an early age and characterized by joint impairment, chronic pain, and reduced quality of life ^5–10^.

Current efforts to prevent the development of HA are mainly focused on management of acute joint bleeding and optimizing prophylactic replacement therapy ^2,11,12^. However, the widespread adoption of the modern hematological care has not conferred considerable protection against the development of HA; even in patients with hemophilia receiving intermediate- and high-dose prophylaxis, many still develop HA ^13^. Pathological changes of HA begin with the first joint bleedings, either breakthrough or sub-clinical, usually during the first years of life ^4,14^. They consist in progressive and irreversible alterations induced by the direct and indirect toxicity of the free blood in the joint space. The joint replacement surgery, such as Total Knee Arthroplasty (TKA), is considered an intervention for alleviating pain and restoring joint function in patients with end-stage HA. However, it is associated with a high incidence of complications, primarily attributed to septic loosening and recurrent postoperative bleeding ^13^. Given no disease modifying therapy available to intervene in the HA perpetuating process, HA remains a persistent problem and challenge for hemophilia patients. Thus, to understand the pathogenesis of HA and subsequently explore possible targets for therapy is necessary.

One distinctive pathophysiological manifestation of HA is cartilage degeneration ^15^. Cartilage is a rather inert tissue, consisting of matrix proteins and only one cell type, chondrocytes that are responsible for production of matrix synthesis and rely on synovial fluid for nutrients as cartilage lacks blood supply ^16,17^. Nevertheless, recurrent intra-articular bleeding creates a ‘toxic’ environment to cartilage, by inducing synovial inflammation, hemosiderin accumulation and excessive mechanical stress ^15,18^. As a consequence, dysregulation of metabolism and abnormal apoptosis occur in chondrocytes, ultimately resulting in deterioration of the cartilage matrix ^2^.

DNA methylation, a core epigenetic mechanism, is high dynamic and susceptible to cues from the environment ^19,20^. DNA methylation is the addition of a methyl group to the cytosine residue of DNA molecules, which is catalyzed by DNA methyl transferase (Dnmt) family of proteins that is comprised of three members: Dnmt1, Dnmt3a, and Dnmt3b. DNA methylation is mainly located in the gene regulatory region, usually repressing gene expression by blocking the transcriptional accessibility of regulatory genomic regions ^21^. Recently, multiple studies confirmed the important role of abnormal DNA methylation in the pathogenesis of arthritis ^22,23^. However, the epigenetic mechanisms underlying HA-related cartilage degradation have not been explored.

In this study, we conducted a genome-wide DNA methylation study with the goal of identifying differentially methylated genes (DMGs) and pathways for HA. Further, we knocked down *Tnxb*, a key DMG, in primary mouse chondrocytes and hemophilia A (*F8^-/-^*) mice. The biological function and underlying mechanisms of TNXB in HA progression were also investigated *in vivo* and *in vitro*. Our findings provide a new mechanism for articular cartilage lesion in HA, which may lead to the discovery of novel therapeutic targets for the treatment of HA.

## Results

### Articular cartilages of HA patients show severe damage and aberrant elevations of Dnmt1 and Dnmt3a

In this study, we collected cartilage samples in tibial plateau from OA and HA patients undergo total knee replacement (TKR) surgery. Consistent with previous reports, the HA cartilage exhibited severe damage and significant hemosiderin deposition, compared with that of OA (Figure 1A). MRI analysis revealed the subchondral bone loss in HA patients (Figure 1B). ABH staining further assessed the cartilage degeneration, and detected a substantial sulfated glycosaminoglycan (sGAG) depletion in HA cartilages (Figure 1, C and D). As expected, comparing to OA, HA cartilages showed prominent decrease in cartilage matrix protein Col2a1 and increase in expression of Mmp13, the primary enzymes responsible for cartilage degeneration (Figure 1, E and F). Additionally, obvious increase in Dnmt1 and Dnmt3a protein levels was also detected in HA cartilages (Figure 1, G and H), whereas no significant changes were found in Dnmt3b (Figure 1-Figure supplement 1). These data confirmed the severe damage in articular cartilage and subchondral bone of HA patients.

**Figure 1.**
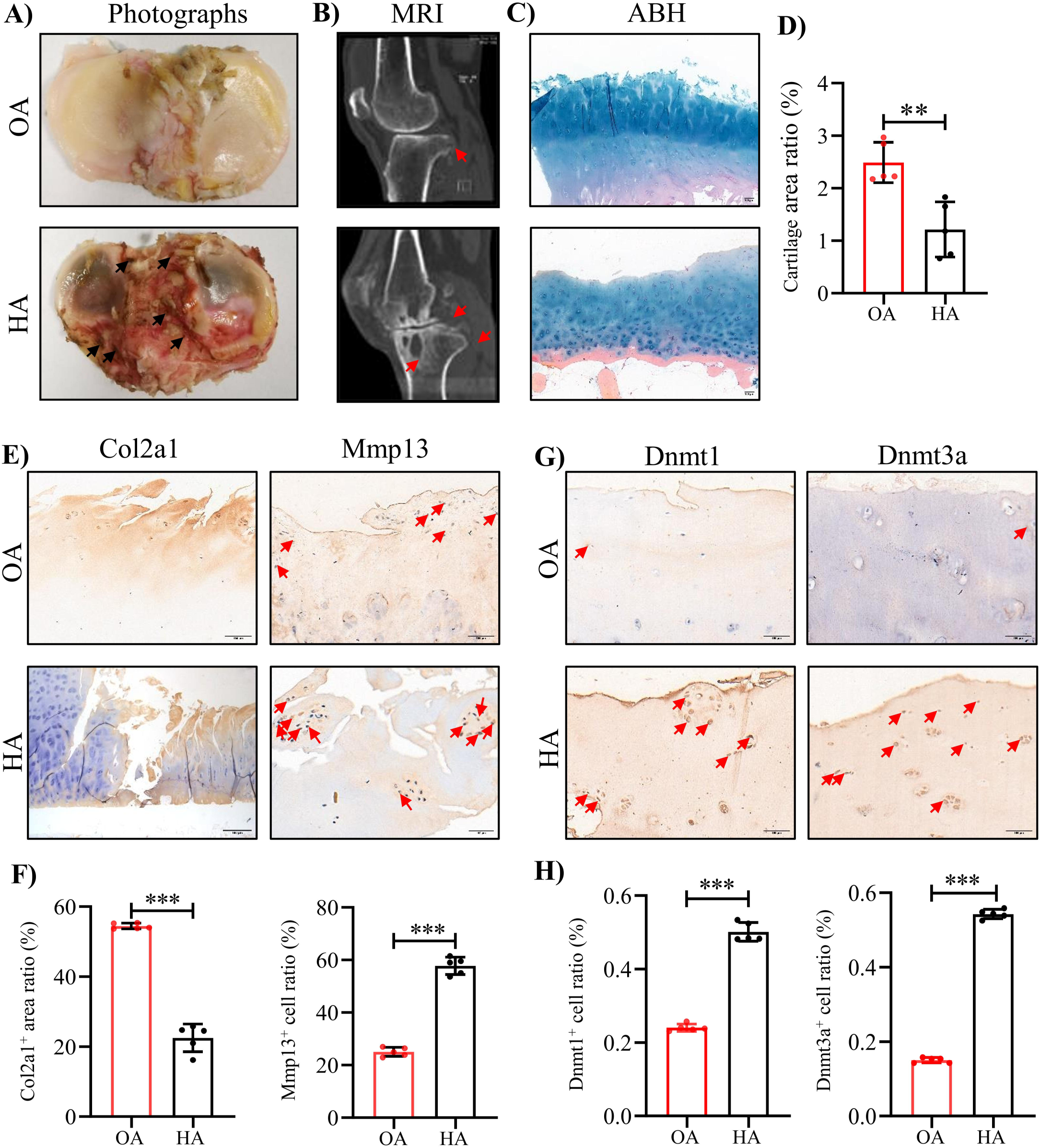
The severe cartilage damage in human HA. **(A)** Human cartilage samples in the tibial plateau were obtained from patients with OA and HA. Black arrow indicates hemorrhagic ferruginous deposits. **(B)** MRI imaging of the knee joint in patients with HA and OA. Red arrow indicates the wear area. **(C)** Representative images of ABH/OG staining for HA and OA cartilage. **(D)** The degree of cartilage degeneration was quantified according to the OARSI score. **(E)** Representative IHC staining of COL2A1 and MMP13. **(F)** Quantification of the proportion of COL2A1 and MMP13 positive regions in human cartilage. **(G)** Representative IHC staining of DNMT1 and DMMT3A. **(H)** Quantification of the proportion of NMT1 and DMMT3A positive cells in human cartilage. Red arrows indicate positive cells. Scale bar: 100 μm. Data were presented as means ± SD; n = 5 per group. And analyzed by 2-tailed unpaired parametric Student’s t test, \*\**P* < 0.01, \*\*\**P* < 0.001.

### Genome-wide DNA methylation analysis reveals the epigenetic landscape of HA cartilage

To understand the methylation profile associated with HA cartilage degeneration, genome-wide DNA methylation alterations were examined in 10 subjects (5 OA cartilages and 5 HA cartilages) by using 850K ChIP. The methylation levels of whole genome-wide CpGs were in a classic bimodal distribution where most CpGs were either slightly or highly methylated (Figure 2A). Of note, global DNA methylation levels exhibited systematic differences between HA and OA, as illustrated by their separation pattern in principle component analysis (Figure 2B). Overall, 1288 significant differentially methylated regions (DMRs) (P<0.05 after the Benjamini & Hochberg correction for multiple testing) were identified, with a mean fold change in methylation difference of 0.90 (Figure 2C, and Supplement file 3a). Compared with the OA, 69.95% and 30.05% DMRs were hypermethylated and hypomethylated in HA, respectively (Figure 2-Figure supplement 1). Given the specificity of DMR, we then determined whether the identified DMRs were preferentially colocalized with some specific genomic features. The large majority (93.49%) of these DMRs were located in gene promoters, 5’-UTRs, 3’-UTRs, and exons, with only 6.51% overlapping intergenic regions (Figure 2-Figure supplement 2).

**Figure 2.**
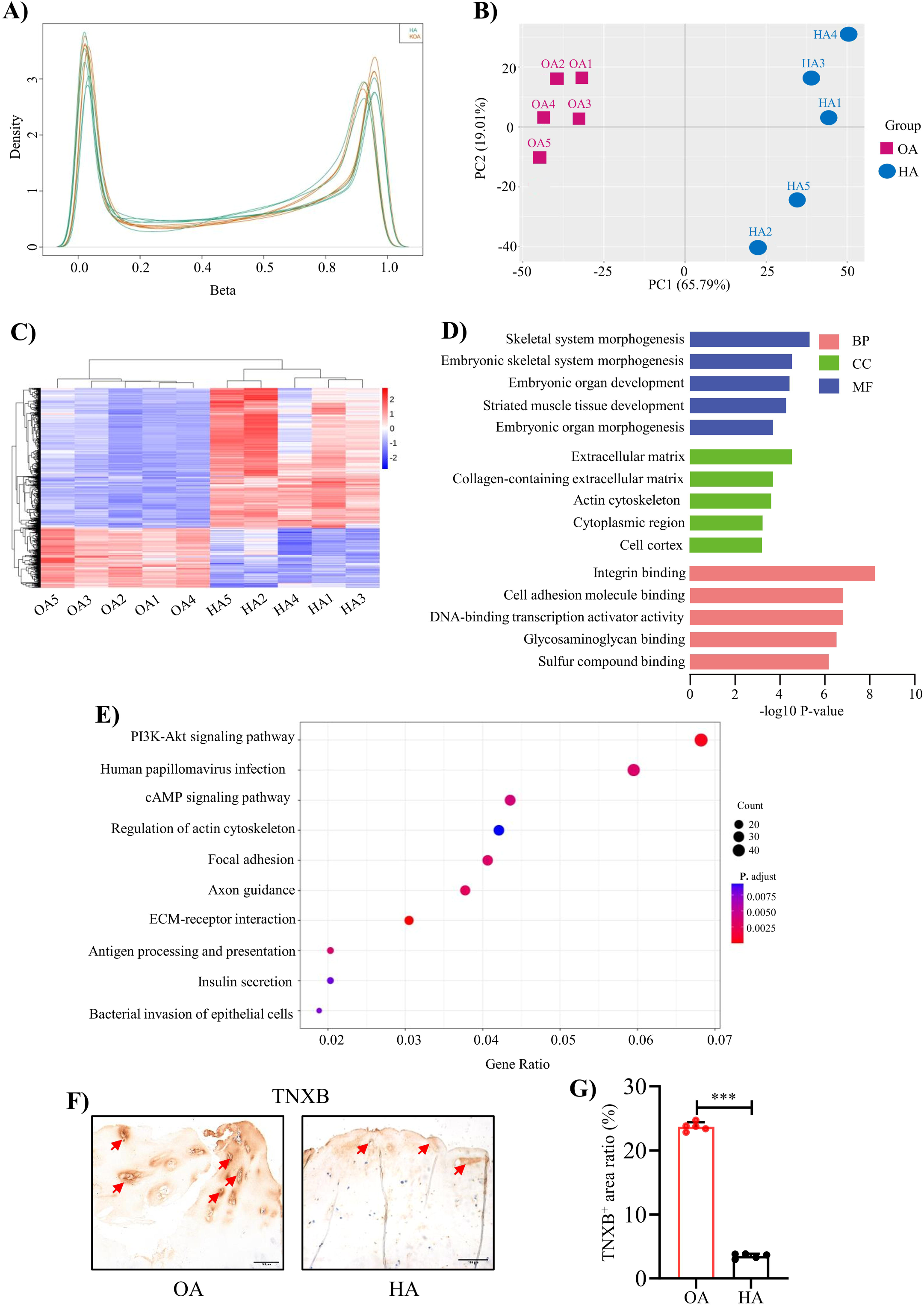
Genome-wide DNA methylation profile in HA cartilage and biological functions of DMR-Related genes. **(A)** Distribution of methylation level density of CpGs. Note: X = degree of methylation; Y = the CpG site density corresponding to the level of methylation. **(B)** Principal component analysis of DNA methylation data. **(C)** Heatmap shows the 700 significant DMRs between HA and OA. **(D)** Enriched GO terms for DMR-related genes. **(E)** KEGG enrichment analysis of DMR-related genes. **(F)** Representative IHC staining of TNXB. Red arrows indicate positive areas. Scale bar: 100 μm. **(G)** Quantification of the proportion of TNXB positive regions in human cartilage. Data were presented as means ± SD; n = 5 per group. And analyzed by 2-tailed unpaired parametric Student’s t test, \*\*\**P* < 0.001.

To determine the biological functions of these significant DMRs, GO and KEGG enrichment analysis were performed on DMR-related genes (DMGs). GO analysis revealed enrichment for terms largely related to extracellular matrix (ECM) organization, skeletal system development (Figure 2D, Supplement file 3b). The top two KEGG pathways were ECM-receptor interaction (*P* value = 5.2×10^-6^) and PI3K-Akt signaling pathway (*P* value = 5.3×10^-4^) (Figure 2E, Supplement file 3c). DMGs presented in these pathways have mostly been proved to play an important role in cartilage matrix homeostasis, such as *COL1A1*, *COL1A2*, *COMP* ^27–29^. Among them, we found TNXB were significantly low expressed in human HA cartilage compared to OA cartilage by using IHC staining (Figure 2, F and G). However, TNXB was not differentially expressed in OA and HA synovial membranes (Figure 2-Figure supplement 3).

### Decreased expression of Tnxb is associated with cartilage degradation by joint bleeding in *F8^-/-^* mice

To assess the potential relevance of TNXB in HA *in vivo*, we established a mouse model for HA, in which *F8^-/-^* mice received a needle puncture injury to the knee joint ^30,31^. This needle puncture induced excessive bleeding and caused severe hemarthrosis (Figure 3-Figure supplement 1). Analysis of cartilage tissue sections by Toluidine blue staining uncovered that joint bleeding remarkably promoted cartilage degeneration at 4 and 8 weeks after injury (Figure 3, A and B). IHC staining further demonstrated lower expression of Col2a1 and increased expression of Mmp13 (Figure 3, C-F). Next, micro-CT was used to assess the influence of hemarthrosis in osteogenesis. An obvious osteopenia was detected in subchondral bone at 4 and 8 weeks post injury (Figure 3-Figure supplement 2). As expected, the above histopathological changes in *F8^-/-^* mice following needle puncture were similar with what we observed in Human HA. Meanwhile, decreased expression of Tnxb and increased expressions of Dnmt1 and Dnmt3a were showed in articular cartilages 4 and 8 weeks following injury (Figure 3, G-L). Above results suggest that the suppression of TNXB in cartilage promotes the HA development.

**Figure 3.**
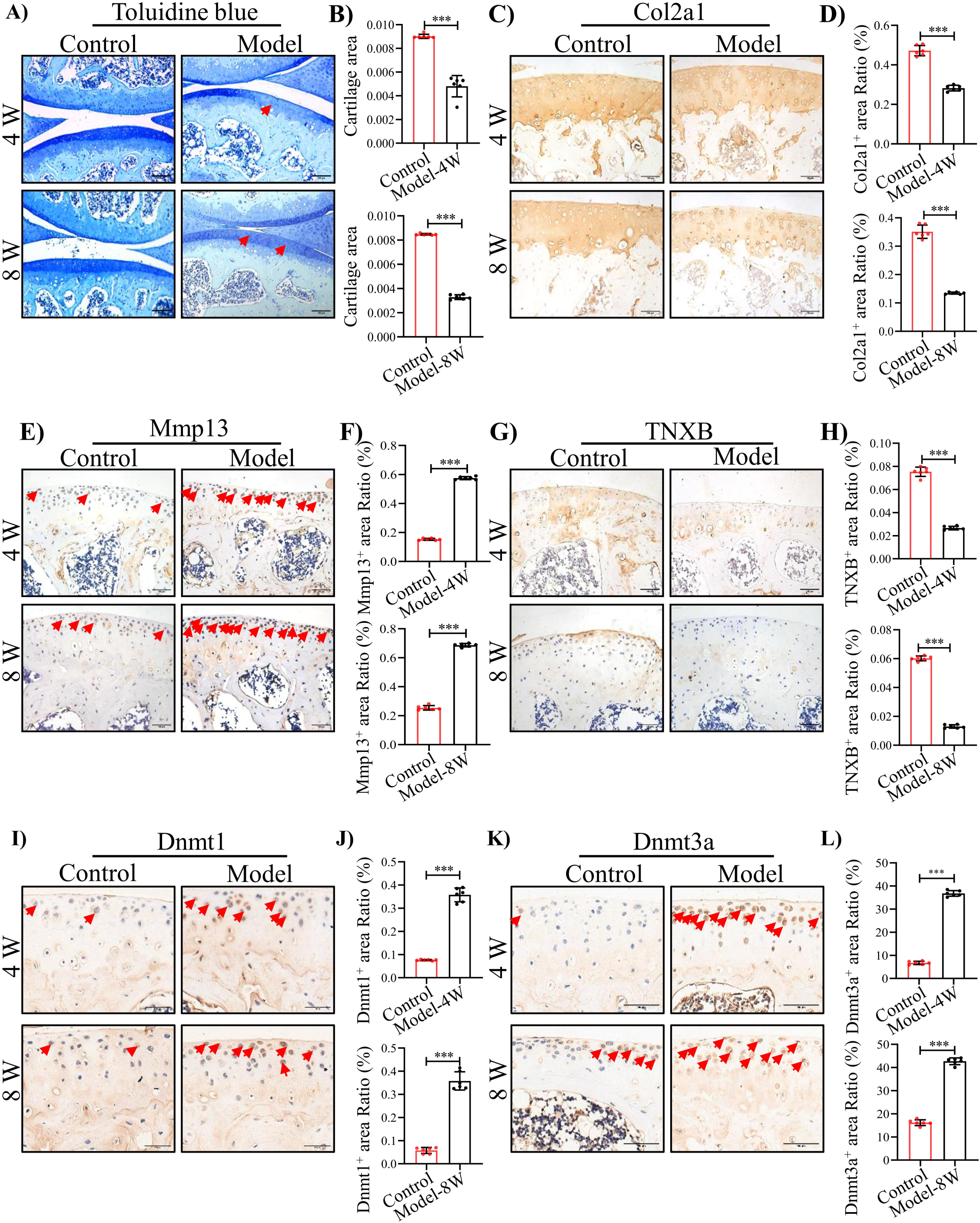
TNXB expression is drastically reduced in HA mouse cartilages. **(A)** Representative images and **(B)** Quantitative analysis of Toluidine blue staining for knee sections of *F8^-/-^* mice at 4 and 8 weeks following injury. Representative IHC staining of **(C)** Col2a1, **(E)** Mmp13, **(G)** Tnxb, **(I)** Dnmt1 and **(K)** Dnmt3a in *F8^-/-^* mice at 4 and 8 weeks post injury. Quantification of the proportion of **(D)** Col2a1, **(F)** Mmp13, **(H)** Tnxb, (**J)** Dnmt1 and (**L)** Dnmt3a positive regions. Scale bar: 100 μm. Data were presented as means ± SD; n = 6 mice per group. And analyzed by 2-tailed unpaired parametric Student’s t test, \*\*\**P* < 0.001.

### *TNXB* knockdown impairs chondrocyte metabolism and aggravates the progression of HA

To verify the relationship between TNXB and methylation levels, we treated primary mouse chondrocytes with DNA Methyltransferase inhibitors RG108 or 5-Aza-dc, well-known to block DNA methylation ^32,33^. After 24 hours treatment, RG108 or 5-Aza-dc (10, 25 μM) significantly up-regulated the mRNA level of *Tnxb*, indicating that the DNA methylation inhibits the expression of TNXB in chondrocytes (Figure 4-Figure supplement 1). To further identify the biological functions of TNXB in HA, we first knocked it down by specific siRNA in primary mouse chondrocytes. The efficiency of *Tnxb* knockdown was confirmed by results of qPCR and Western blot experiments (Figure 4, A-C). When compared with the siRNA-control group, we found that *Tnxb* knockdown led to a decrease in *Col2a1* mRNA expression and an increase in *Mmp13* mRNA expression (Figure 4, D and E). Western blot and immunofluorescence staining analysis also showed a downregulated expression of Col2a1 protein and upregulated expression of Mmp13 protein (Figure 4, E-H). Those observations revealed that TNXB positively regulated the metabolism of chondrocyte ECM.

**Figure 4.**
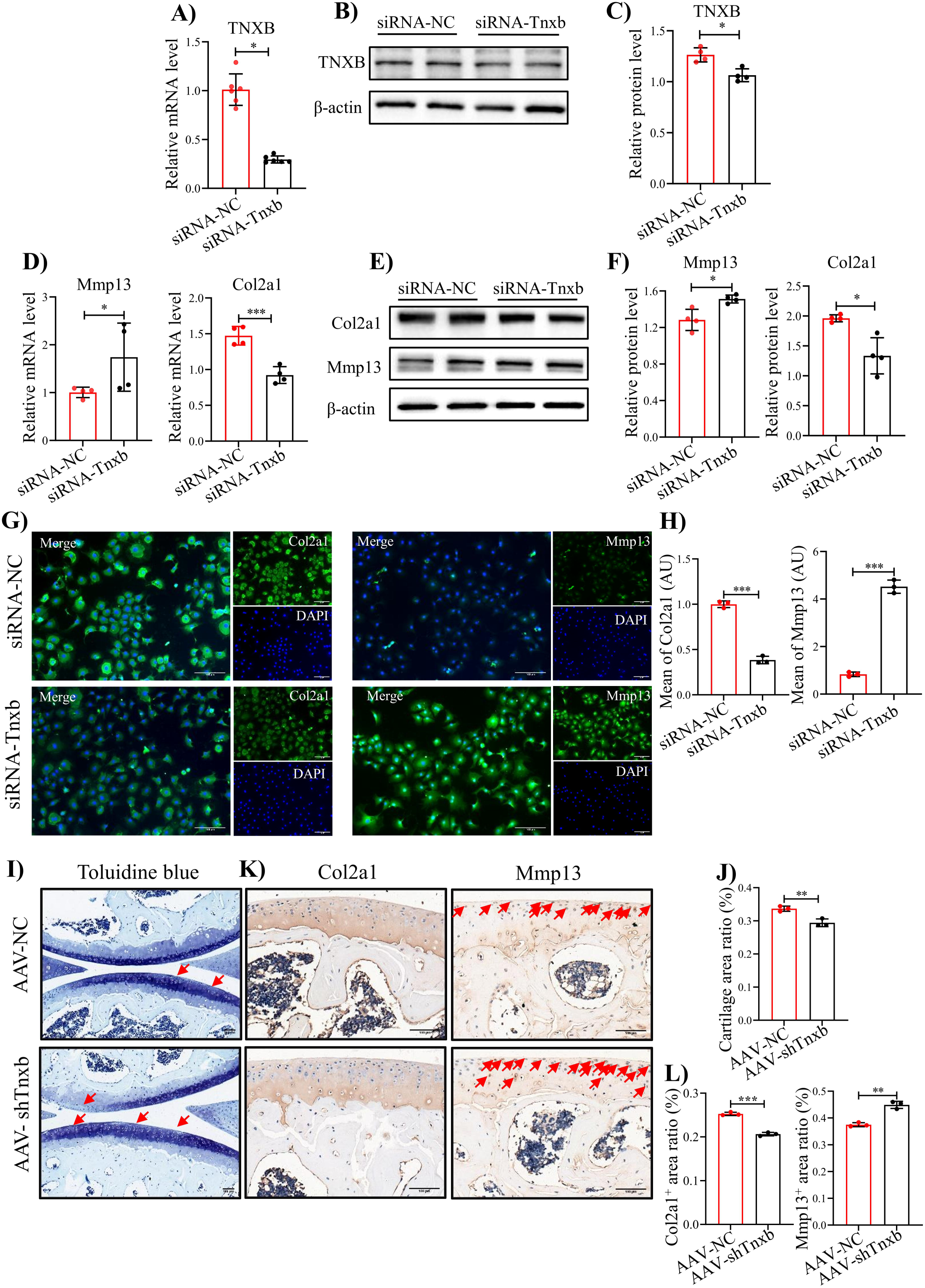
knockdown of Tnxb in chondrocytes induces extracellular matrix degradation and accelerates the progression of HA. **(A)** qPCR and **(B)** Western blotting analysis for Tnxb in mouse chondrocytes treated with siRNA-NC or siRNA-*Tnxb*. **(C)** Corresponding quantification analysis of Tnxb protein. Quantification of mRNA levels for **(D)** *Mmp13* and*Col2a1* in *Tnxb*-KD chondrocytes. **(E)** Western blot and **(F)** corresponding quantification analysis for Mmp13 and Col2a1 protein. **(G)** Representative immunofluorescence images of Col2a1 and Mmp13 expression in *Tnxb*-KD chondrocytes. **(H)** Quantification of Col2a1 and Mmp13 fluorescence intensity. Representative images of **(I)** Toluidine blue staining and **(K)** IHC staining of Mmp13 and Col2a1 for knee sections of *F8^-/-^* mice at 4 weeks after Intra-articular injection of AAV-sh*Tnxb*. **(J)** Quantitative detection of the area of the tibial cartilage area. **(L)** Quantification of the proportion of Col2a1 and Mmp13 positive regions. Red arrow indicates the wear area. Scale bar: 100 μm. Data were presented as means ± SD; n ≥ 3 in each group. And analyzed by 2-tailed unpaired parametric Student’s t test, \**P* < 0.05, \*\**P* < 0.01, \*\*\**P* < 0.001.

Subsequently, we explored the effects of TNXB on HA cartilage degradation *in vivo*, using intra-articular injection of adeno-associated virus (AAV) carrying *Tnxb*-specific short hairpin RNA (shRNA) in *F8^-/-^* mice. 4 weeks after injection, more severe cartilage degeneration was observed in *Tnxb*-KD mice, compared with vehicle group (Figure 4, I and J). In addition, *Tnxb*-KD mice showed decreased expression of Col2a1 and increased expression of Mmp13 in cartilages (Figure 4, K and L). The subchondral bone of *Tnxb*-KD mice was significantly less than vehicle group mice (Figure 4-Figure supplement 2). Collectively, the above results implied that TNXB knockdown in cartilage accelerated the HA development.

### *TNXB* deficiency sensitizes chondrocytes to apoptosis by inhibiting AKT activation

Since blood exposure in joint cavity strongly causes chondrocyte apoptosis (Figure 5-Figure supplement 1), we next explored the functional mechanisms of TNXB in HA chondrocyte apoptosis. *Tnxb* knockdown induced the apoptosis in chondrocytes, as evidenced by an increased proportion of Tunel-positive cells (Figure 5, A and B). Meanwhile, protein expression levels of pro-apoptotic Bax and Cleaved-Caspase3 were significantly increased in *Tnxb*-KD chondrocytes, while anti-apoptotic Bcl-2 was decreased (Figure 5, C-F). Moreover, AAV-shTnxb injection also increased the number of Tunel-positive cells in articular cartilages in HA mice (Figure 5, G and H). IHC staining confirmed that Bcl-2 expression was significantly reduced in *Tnxb*-KD HA mice accompanied by enhanced expressions of Bax and Caspase-3 (Figure 5, I-L).

**Figure 5.**
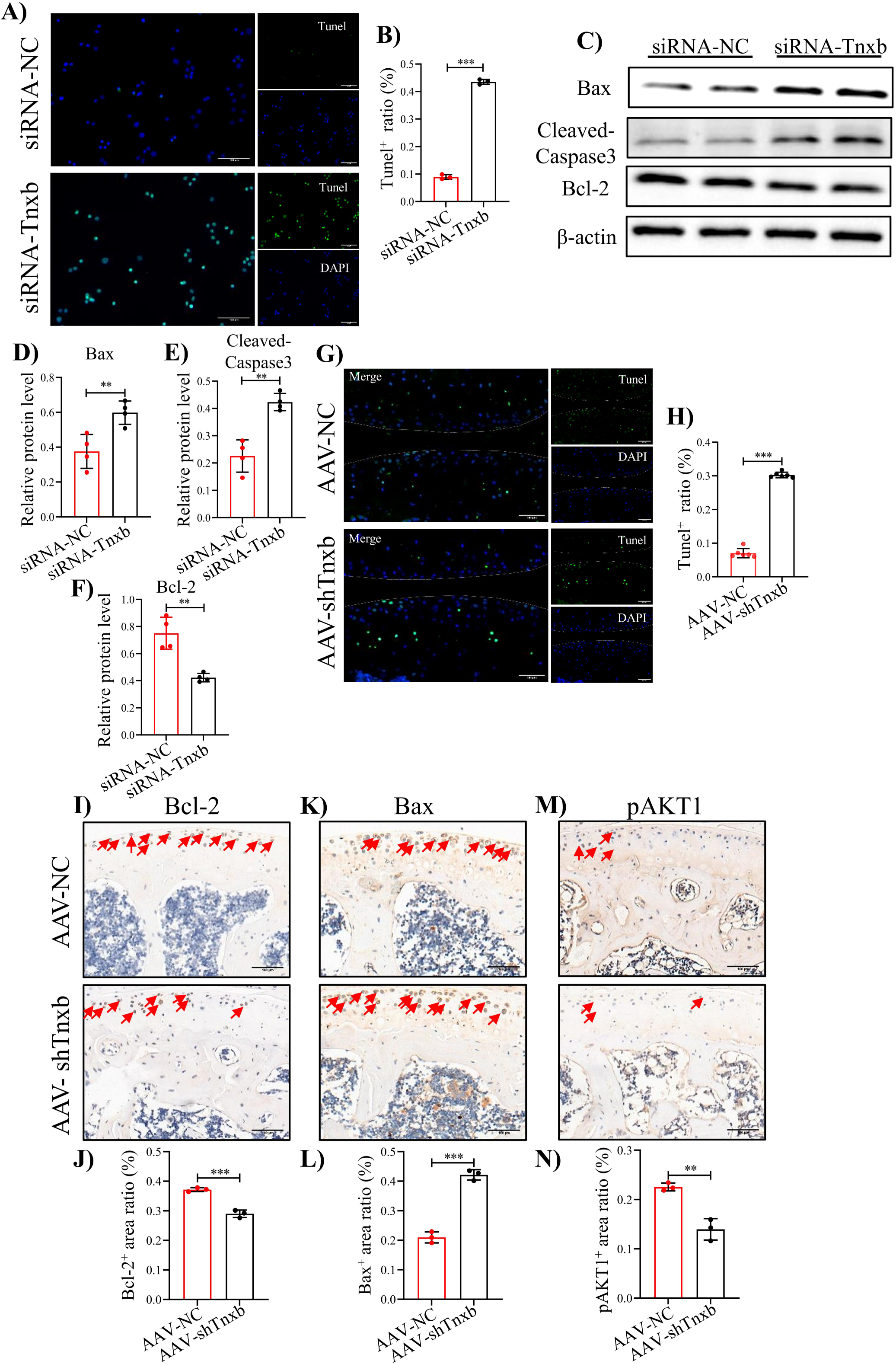
Tnxb knockdown increases apoptosis in chondrocytes and HA cartilages. **(A)** Representative images of TUNEL staining in Tnxb-KD chondrocytes **(B)** Quantification for TUNEL positive cells. **(C-F)** Western blot and corresponding quantification analysis for Bax, Cleaved-Caspase3, and Bcl-2. **(G)** TUNEL staining for apoptosis in articular cartilage from Tnxb-KD HA mice. **(H)** Quantification for TUNEL positive cells in articular cartilage. Representative IHC staining of Bax **(I)**, Bcl-2 **(K)**, and p-Akt1 **(M)** in articular cartilage from *Tnxb*-KD HA mice. Corresponding quantification of the proportion of **(J)** Bax, **(L)** Bcl-2 and **(N)** p-Akt1 positive regions. Red arrows indicate positive cells. Scale bar: 100 μm. Data were presented as means ± SD; n ≥ 3 in each group. And analyzed by 2-tailed unpaired parametric Student’s t test, \*\**P* < 0.01, \*\*\**P* < 0.001.

It is well known that the PI3K/Akt signaling pathway regulates apoptosis ^34–36^, and our KEGG analysis and IHC staining confirmed the involvement of this pathway in HA cartilage degeneration (Figure 5, M and N). We, therefore, analyzed PI3K/Akt pathway following *Tnxb* knockdown. In *Tnxb*-KD chondrocytes, p-Akt protein level decreased significantly, while Akt level remained unchange (Figure 6, A and B). Next, we sought to investigate whether Akt activation could attenuate chondrocyte apoptosis induced by *Tnxb* knockdown. AKT agonist SC79 was used to treated *Tnxb*-KD chondrocytes, and then its efficiency was verified by western blotting (Figure 6-Figure supplement 1). 24 hours after SC79 treatment, the expressions of Bcl-2 and Col2a1 (Figure 6, A, B and F) were upregulated, while Bax, Mmp13 and Cleaved-Caspase9 expressions were markedly downregulated in *Tnxb*-KD chondrocytes (Figure 6, A, C, E and G). These findings indicate that *TNXB* knockdown induced chondrocyte apoptosis and ECM degradation through regulating AKT activation.

**Figure 6.**
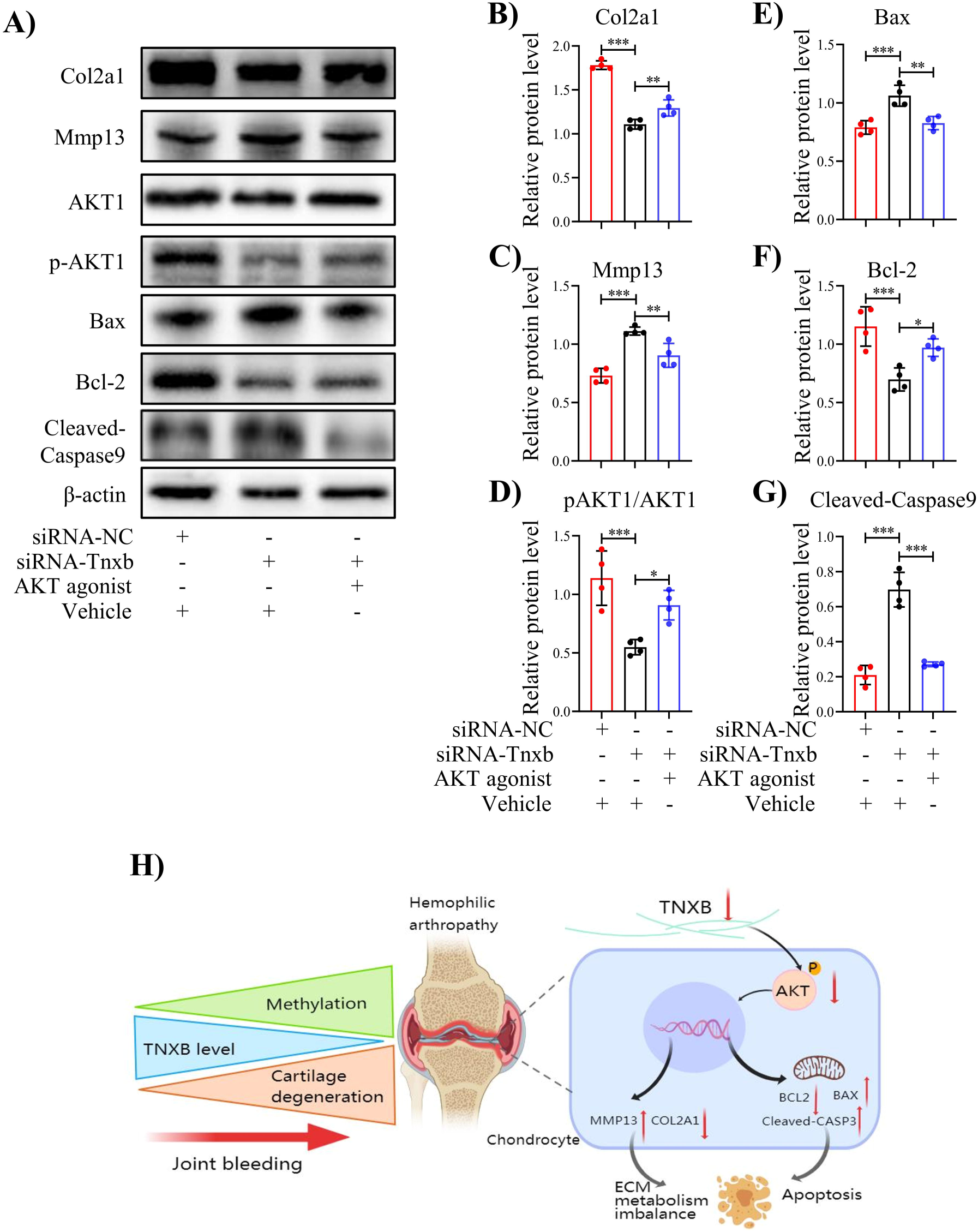
Treatment with AKT agonist decreases apoptosis in TNXB-KD chondrocytes. **(A)** Western blot analysis for Col2a1, Mmp13, Bax, Bcl-2, p-Akt1, Akt1 and Cleaved-Caspase9 in Tnxb-KD chondrocytes treated with SC79 (0 or 10μM) for 24 hours. **(B-G)** Corresponding quantification of these proteins. **(H)** Schematic of the role of TNXB in the pathogenesis of HA cartilage degeneration. Data were presented as means ± SD; n = 4 per group. And analyzed by 2-tailed unpaired parametric Student’s t test or one-way ANOVA with Tukey’s multiple comparisons test, \**P* < 0.05, \*\**P* < 0.01, \*\*\**P* < 0.001.

## Discussion

HA, a common manifestation of hemophilia, is typical characterized by dramatic cartilage degradation ^6^. Thus, the goal of our work was to provide a novel insight into pathogenic mechanisms underlying HA cartilage degeneration. Figure 6 H summarized our findings. Genome-wide DNA methylation analysis revealed the DNA methylation changes in cartilages from hemophilia patients and identified *Tnxb* gene associated with HA. The decreased expression of TNXB protein was also confirmed in both human and mouse HA cartilages. Further, *Tnxb* knockdown promoted ECM catabolism and chondrocyte apoptosis, and eventually accelerated the development of HA in *F8^-/-^* mice. Meanwhile, decreased TNXB expression inhibited the activation of AKT *in vivo* and *in vitro*. Notably, AKT agonist enhanced ECM synthesis and suppressed apoptosis in *Tnxb*-KD chondrocytes. From these findings, it could be hypothesized that abnormal methylation and decreased expression of TNXB are responsible for disruption of cartilage homeostasis in HA.

In hemophlia patients, recurrent joint bleeding creates a toxic environment including multifaceted processes such as hemosiderin deposition, inflammatory response and mechanical pressure overload ^37–40^. DNA methylation is a dynamic epigenetic modification in response to environment. Since there are fewer epigenetic studies on the pathological mechanisms of HA, we performed genome-wide DNA methylation analysis, we found 1288 significant DMRs associated with HA with 69.95% hypermethylated, which was also supported by a corresponding increase in Dnmt1 and Dnmt3a protein levels in HA cartilages. These results are similar to previous reports showing that aberrant elevations of Dnmt1 and Dnmt3a promote cartilage degeneration in destabilization of medial meniscus and aging models ^33,41^.

DNA methylation at gene promoter region is one of the important epigenetic mechanisms in the regulation of gene expression ^42,43^. The identified HA-related DMRs showed high proportion in transcriptional start sites. These data suggest that DNA methylation is a critical mediator in the pathology of HA. Nevertheless, due to the severe erosion of HA cartilage, the mRNA levels of DMR-related genes were not explored in this study. Besides, our sample size for DNA methylation sequencing is relatively small. Thus, further studies using additional more HA samples will be conducted to investigate the possible transcriptional alteration of identified genes as well as their potential roles in HA development.

Furthermore, our enrichment analysis showed that DMR-related genes were significantly enriched in ECM organization and ECM-receptor interaction terms. It is consistent with previous reports that ECM acts as an epigenetic informational entity capable of transducing and integrating intracellular signals via ECM-receptor interaction ^44–46^. DMR-related genes enriched in these terms, many are known to be critical compositions of cartilage matrix, such as *COMP*, *COL1A1* and *COL1A2* ^27–29^. In Articular cartilage, ECM provides nutrients and mechanical support for chondrocytes ^47^, and its age-related stiffening epigenetically regulates gene expression and compromises chondrocyte integrity ^41^. Above findings highlight that ECM-related genes are potential candidates for HA.

Tenascin-X (TNXB), an ECM glycoprotein, is the top differentially methylated gene in our DNA methylation analysis of HA cartilages. In addition, DNA methylation of TNXB has also been reported in whole blood from rheumatoid arthritis patients and in retinal pigment epithelium from patients with age-related macular degeneration ^48,49^. Using IHC staining, we detected the downregulated expression of TNXB in human and mouse HA cartilages. TNXB belongs to tenascin family, whose members (TNC, TNR, TNXB, and TNW) share a similar domain pattern: an N-terminal oligomerization domain, a series of epidermal growth factor-like repeats, a variable number of fibronectin-type III (FNIII) repeats and a C-terminal fibrinogen-like domain ^50^. Emerging evidence suggests the involvement of TNC in cartilage development and degeneration in arthritis ^51,52^. Here, we prove for the first time a role of TNXB in cartilage homeostasis *in vivo* and *in vitro*. Specifically, *Tnxb* knockdown contributed to an increase in Mmp13 expression and decrease in Col2a1 expression in chondrocytes. In agreement with these findings, knockdown of *Tnxb* in cartilages of *F8^-/-^* mice resulted in accelerated HA-like lesions. It has been previously reported that TNXB blocks the interaction between TGF-β and its receptor in endothelial cells. Since a potent role of TGF-β in chondrocyte differentiation, we examined the effect of TNXB on TGF-β signaling, but did not observe significant alterations in this signaling in *Tnxb*-KD chondrocytes (Figure 6-Figure supplement 2, A and B). The FNIII repeats domain of TNXB undergoes alternative splicing to interact with different ECM proteins and growth factors, and then affects ECM network formation and three-dimensional collagen matrix stiffness ^53^. Thus, future study would benefit from exploring the effect of the FNIII repeats domain in HA development to further clarify the specific mechanisms of TNXB.

The process of HA development is accompanied by the chondrocyte apoptosis which in part promotes the ECM degeneration ^15,54^. We also observed remarkably increased apoptosis following joint bleeding in *F8^-/-^*mice. Interestingly, *Tnxb* knockdown could significantly induce apoptosis in chondrocytes *in vitro*, and even aggravate apoptosis in mouse HA cartilage, by enhancing expression of markers associated pro-apoptosis (Bax, C-Caspase3) and suppressing anti-apoptotic proteins (Bcl-2). In TNXB family, TNC was reported to inhibit human chondrosarcoma cell apoptosis by activation of AKT ^55^. PI3K signaling pathway regulated AKT phosphorylation to suppress apoptosis ^56,57^. Our KEGG analysis of DMGs showed the potential association of PI3K/AKT signaling with HA. We then detected much lower phosphorylation level of AKT in *Tnxb*-KD chondrocytes and HA models. These results are in line with previous report showing that the expression of p-AKT1 was down-regulated in the articular cartilage of patients with HA compared with that in patients with OA ^58^. Notably, we further demonstrated that AKT agonist effectively restored the abnormal Mmp13 expression and apoptosis induced by *Tnxb* knockdown. Recent studies have shown that AKT signaling affected cartilage metabolism by regulating Mmp13 expression ^59,60^. These findings indicate that AKT1 activation mediates the effect of TNXB on HA chondrocytes apoptosis and ECM catabolism. Thus, therapeutic effect of AKT agonist on HA development deserves further study.

## Conclusions

Conclusively, our study provides the first comprehensive methylation profile in the progression of knee HA, and shows that abnormal expression of TNXB is important in HA cartilage degeneration by suppressing the activation of AKT1. These findings provide a new insight into mechanisms responsible for HA disease and suggest TNXB as a potentially novel therapeutic target for HA treatment.

## Materials and Methods

### Cartilage specimen collection

Osteoarthritis (OA) is often used as “disease” control to reveal the characteristics in HA ^24,25^, although the mechanistic and phenotypic are different between HA and OA. In this study, 5 HA and 5 OA knee cartilage specimens were collected from patients undergoing total knee replacement surgery. Knee cartilage tissues were dissected and then rapidly frozen in liquid nitrogen or fixed in 4% paraformaldehyde. All subjects were Chinese Han and their demographic characteristics were listed in Supplement file 1. The human cartilage samples were obtained from Department of Orthopedic Surgery at The First Affiliated Hospital of Zhejiang Chinese Medical University. This study was approved by the Ethics Committee of the First Affiliated Hospital of Zhejiang Chinese Medical University (2019-ZX-004-02).

### Magnetic Resonance Image (MRI)

MRI examination: a GE 3.0 T Signa Excite superconducting MRI machine with a 32-channel body coil, a gradient field of 24 mT/m, and a creep rate of mT/(m.s) as well as a Siemens Verio 3.0 T superconducting magnetic resonance imaging machine with a 16-channel phased array coil were applied. An ultrashort echo time (UTE) pulse sequence (echo time 0.07 ms) was performed in the sagittal plane of the knee joint of the affected limb. Scanning parameters: layer thickness of 0.8 mm, field of view of 320 mm×320 mm, matrix of 160 mm×160 mm, intra-layer resolution of 0.6 mm×0.6 mm, and excitation of 2 times. Scanning software: NooPhase Wrap, Variable Bandwidth, Tailored RF.

### Genome-wide DNA methylation profiling

Genomic DNA was prepared from cartilage specimens using QIAamp DNA Blood Mini Kit (QIAGEN, Germany) according to the manufacturer’s instructions and was stored at −80 °C until used. Then bisulfite treatment of genomic DNA was performed with the EZ DNA methylation kit (Zymo Research) according to manufacturer’s procedure. Genome-wide DNA methylation patterns were evaluated by Infinium Human Methylation 850 K BeadChips (Illumina), which determine the methylation levels of 853,307 CpG sites. R/Bioconductor (version 3.3.3) package ChAMP was used to process the Illumina intensity data (IDAT) files from the chip. DNA methylation level of each CpG site was described as a β value, ranging from zero (representing fully unmethylated) to one (representing fully methylated). Probes that had a detection *P* value of > 0.01 and those located on the X and Y chromosome were filtered away. We also removed SNP-related probes and all multi-hit probes. BMIQ (Beta MIxture Quantile dilation) algorithm was used to correct for the Infinium type I and type II probe bias. Furthermore, the regression models were adjusted for age and sex. Differentially methylated probes (DMPs) at significance of *P* < 0.05 after the Benjamini & Hochberg correction for multiple testing. Differentially Methylated Regions (DMRs) were identified by Bumphunter with default settings. Additionally, we annotated the DMR-related genes (DMGs), and then performed Gene Ontology (GO) and Kyoto Encyclopedia of Genes (KEGG) pathway analysis by using the online tool DAVID (http://david.abcc.ncifcrf.gov/). The significant pathway/ GO term was identified with *P* value < 0.05.

### Mouse

The FVIII target gene knockout hemophilia (*F8^-/-^*) mouse were purchased from the Shanghai Laboratory Animal Center of the Chinese Academy of Sciences. All mouse were kept in the Laboratory Animal Center of Zhejiang University of Chinese Medicine. The animals were placed in the controlled environmental condition with relatively constant air humidity and temperature as well as manual control of day and night replacement for 12h. The experiment was approved by the Experimental Animal Ethics Committee of Zhejiang University of Traditional Chinese Medicine (LZ12H27001).

### The mouse experimental model of HA

The HA model is constructed according to the description of Hakobyan et al ^26^. *F8^-/-^* male mouse were induced with 3% isoflurane, and 1% isoflurane was maintained under anesthesia. The right knee joint capsule of mouse was pierced with a 30g needle for bleeding modeling, and the left knee of each animal was used as a control group. Mouse were sacrificed at 4 and 8 weeks after injury, and knee joints were collected for further experiments.

### Intra-articular injection

*F8^-/-^* male mouse were induced with 3% isoflurane, and 1% isoflurane was maintained under anesthesia. The right knee joint capsule of mouse was pierced with a 30g needle for bleeding modeling. The needle of the insulin syringe was sagittally inserted into the intercondylar area of the mouse right knee joint, and one dose of 10 μL AAV-Tnxb (GenePharma, concentration: 1×10^10^/10μL) or AAV-NC was injected. After 4 weeks, mouse was sacrificed, respectively, and the right knee joints were collected for further experiments.

### Micro-computed tomography (**μ**CT) analysis

Micro-computed tomography (μCT) (Skyscan 1176, Bruker μCT, Kontich, Gelgium) was used to analyze the knee joints. The area between the proximal tibia growth plate and the tibial plateau was chosen as the region of interest. The parameters collected form μCT were percent bone volume (BV/TV, %), Trabecular thickness (Tb.Th, mm), Trabecular number (Tb.N, 1/mm) and Trabecular Spacing(Tb.Sp, mm).

### Histological analysis

The human cartilage samples were successively fixed in 4% paraformaldehyde for 5 days, decalcified with 14% EDTA solution for 3 months, and the mouse samples were successively fixed in 4% paraformaldehyde for 3 days, decalcified with 14% EDTA solution for 14 days. These samples subsequently embedded in paraffin. Then 3-μm thick sections at the medial compartment of the joints were cut for Alcian blue hematoxylin/Orange G (ABH) or Toluidine Blue staining to analyze the gross cartilage structural changes. Histomorphometry analysis was performed through Osteomeasure software (Decatur, GA).

### Immunohistochemistry

The immunohistochemistry was examined to observe the expressions of protein in cartilage. Briefly, the deparaffinized sections were soaked in 0.3% hydrogen peroxide to block the activity of endogenous peroxidase, then blocked with normal goat serum (diluted 1:20) for 20 min at room temperature. Subsequently, the primary antibodies were added and incubated overnight at 4° C. The next day, the sections were treated with secondary antibodies for 30 min and positive staining was visible by using diaminobenzidine solution (Invitrogen, MD, United States). Then counterstaining was performed with hematoxylin for 5 s. Anti-Col2a1 (Abcam, ab34712, 1:200), Anti-Mmp13 (Abcam, ab39012, 1:300), Anti-TNXB (Proteintech, 13595-1-AP, 1:50), Anti-Bax (Huabio, ET1603-34, 1:200), Anti-Bcl-2 (Huabio, ET1702-53, 1:200), Anti-p-Akt1 (Abclonal, AP0140, 1:200), Dnmt1(Abcam, 1: 200), Dnmt3a (Abcam, 1: 200), Dnmt3b (Huabio, 1: 200) were used in this study.

### Cell isolation and culture

Primary mouse chondrocytes were obtained from the femoral head of 2-week-old C57BL/6 J mouse purchased from the Experimental Animal Center of Zhejiang Chinese Medical University. Mouse were sacrificed and disinfected with 75% ethyl alcohol. Specimens were isolated and rinsed by Phosphate Buffer Saline (PBS) 3 times. Then the cartilage tissues were digested with 0.25% collagenase P at 37 °C for 4h. Chondrocytes were cultured in DMEM/F-12 medium containing 10% fetal bovine serum (FBS) and 1% streptomycin/penicillin in 5% CO_2_ at 37 °C for further experiment.

### Tunel assay

To evaluate the apoptotic cells in the chondrocytes, we performed a Tunel assay according to the manufacturer’s guideline (Beyotime, catalog C1088). Briefly, following deparaffinage and rehydration, sections were permeabilized with DNase-free Proteinase K (20 μg/mL) for 15 minutes at 37°C. Subsequently, slides were treated with Tunel solution and incubated at 37°C for 1 hour in a dark environment and counterstained with DAPI (1:1000; Solarbio) for 10 minutes. Finally, TUNEL positive cells were detected by fluorescence microscope.

### Transfection of siRNA-*Tnxb*

Chondrocytes were transiently transfected with siRNA targeting *Tnxb* (GenePharma) or negative□ control siRNA (scrambled; GenePharma) using X□tremeGENE siRNA transfection reagent (Sigma) following the manufacturer’s instructions. All experiments were performed 72 hours after transfection, and the most effective single siRNA was used for further experiments. After transfection for 48 h, cells were treated with various concentrations of SC79 (10μM, Selleck) for 24 h.

### Cell immunofluorescence

The cells were fixed with 4% paraformaldehyde and blocked with goat serum for 1 hour. Then, cells were incubated with Col2a1 antibody (1:200; Abcam) and Mmp13 antibody (1:100; Abcam) overnight. The next day, the knee joints samples and Chondrocytes with goat antirabbit antibody (1:1000; Invitrogen) conjugated with Alexa Fluor 488 for 1 hour. Cells were stained with DAPI (1:1000; Solarbio).

### Western blot analysis

Total proteins were extracted by RIPA buffer, and concentration of protein was determined by bicinchoninic acid (BCA, Beyotime, Shanghai, China). Proteins were separated on 10% SDS-PAGE gels and transferred to PVDF membranes. The membranes were then blocked with 5% skim milk for 1 h, and were incubated with primary antibodies (Col2a1, Abclonal, A1560, 1:1500), (Mmp13, Abclonal, A11755, 1:1000), (TNXB, Abclonal, A2535, 1:1000), (p-Akt1, Abclonal, AP0140, 1:1000), (Akt1, Huabio, ET1609-47, 1:1000), (Bcl-2, Huabio, ET1702-53, 1:1000), (Bax, Huabio, ET1603-34, 1:1000), (active-pro caspase-3, Huabio, ET1608-64, 1:1000), (Cleaved-Caspase9, Abclonal, A22672, 1:1000), (pSmad2, CST, 8828, 1:1000) overnight at 4°C. The membranes were washed 3 times, and then cultured with the secondary antibody for 1 h. The immunoreactivity was detected with the ECL substrate (Thermo Fisher Scientific, United States) on an Image Quant LAS 4000 (EG, United States). The grey value was calculated by the software of Image J. *β-actin* was used as an internal control in all western blot analysis.

### Quantitative RT-PCR

TRIzol reagent (Invitrogen, United States) was used according to the manufacturer’s protocol to extract total RNA from chondrocytes. The total RNA quantity and purity was evaluated by using NanoDrop 2000 (Thermo Fisher Scientific, United States). Reverse transcription was carried out with cDNA Synthesis Kit (Bmake, Beijing, China). The quantitative real-time-polymerase chain reaction (qPCR) was conducted with SYBR Premix Ex Taq™ II (Takara, Dalian, China), and performed on on a QuantStudio™ 7 Flex Real□ Time PCR System. All of the PCR reactions were repeated 3 times for each gene. The data were analyzed by the 2−ΔΔCT method using *Actb* as the internal control. Supplement file 2 showed the primer sequences of target genes used in the current study.

### Treatment of DNA Methyltransferase inhibitors

To investigate whether DNA methylation affects the expression of TNXB, we treated primary mouse chondrocytes with RG108 or 5-Aza-dc (0, 1, 10, and 25 μM) (RG-108, Beyotime, SD-1137; 5-Aza-2’-deoxycytidine, Beyotime, SD-1047) for 24 h. The cells were then collected for qPCR to detect the mRNA level of *Tnxb*.

### Statistics

All experimental data of this subject were analyzed by GraphPad Prism software version 8.0, and the statistical results of measurement data were expressed as mean ± standard deviation. 2-tailed unpaired parametric Student’s t test or nonparametric Mann-Whitney U test or one-way ANOVA with Tukey’s multiple comparisons test was used for comparison between different groups. The difference was statistically significant with *P* < 0.05.

## Supporting information

Response to Public Reviews

Western blot for source data

## List of abbreviations

HA: hemophilic arthropathy
DMRs: differentially methylated regions
DMGs: DMR genes
TNXB: Tenascin XB
DNMT: DNA methyl transferase
ABH: Alcian blue hematoxylin/Orange G
ECM: extracellular matrix
AAV: adeno-associated virus.

## Study approval

In the current study, the human cartilage samples were obtained from Department of Orthopedic Surgery at The First Affiliated Hospital of Zhejiang Chinese Medical University. This study was approved by the Ethics Committee of the First Affiliated Hospital of Zhejiang Chinese Medical University (2019-ZX-004-02). And all the animal operating procedures in this study were approved by the Experimental Animal Ethics Committee of Zhejiang Chinese Medical University (LZ12H27001).

## Data Availability Statement

The data used to provide support for the results of this study can be obtained from the corresponding authors.

## Availability of data and materials

The datasets used and/or analyzed during the current study are available from the corresponding author on reasonable request.

## Acknowledgments

We appreciate the great help from the Public Platform of Medical Research Center, Academy of Chinese Medical Science, Zhejiang Chinese Medical University.

---

**Figure 1-Figure supplement 1.**
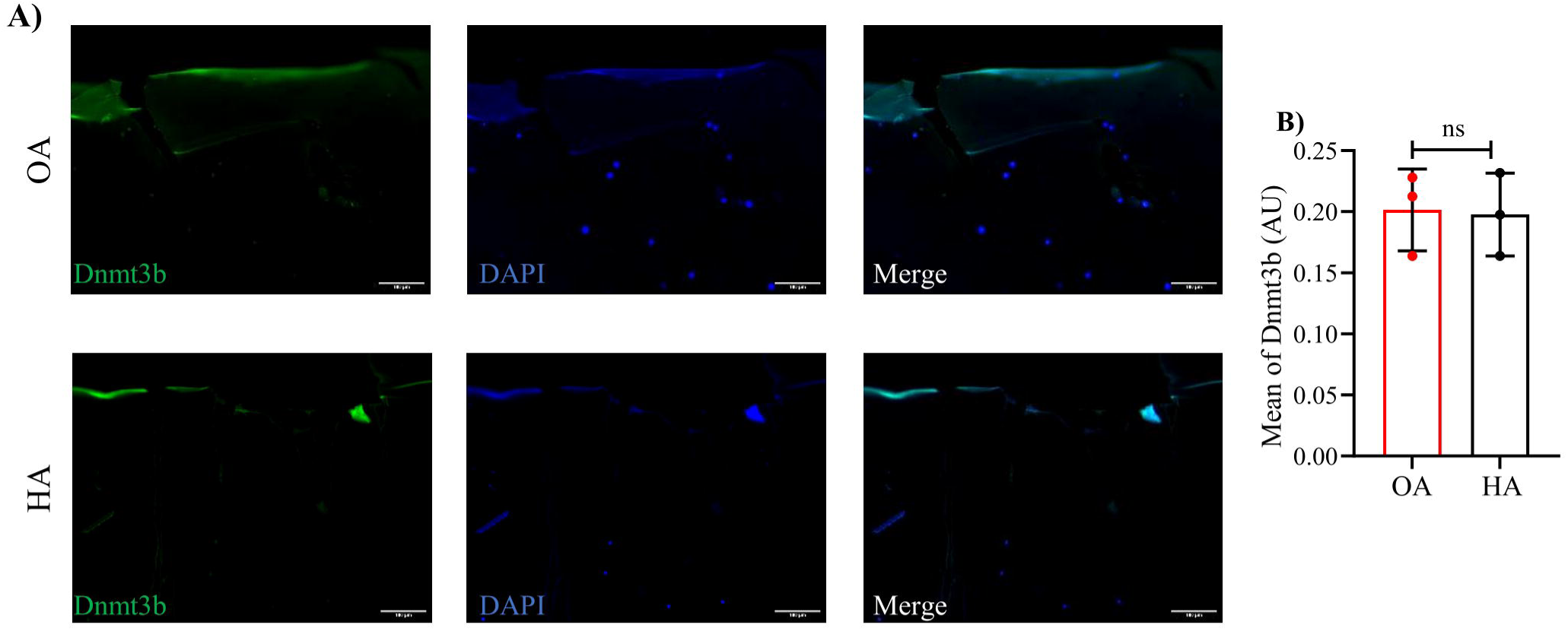
Representative immunofluorescence images (A) and quantification (B) of Dnmt3b in human HA and osteoarthritis cartilages. Scale bar: 100 μm. Data were presented as means ± SD; n = 5 in each group. ns = no significance by unpaired Student’s t test.

**Figure 2-Figure supplement 1.**
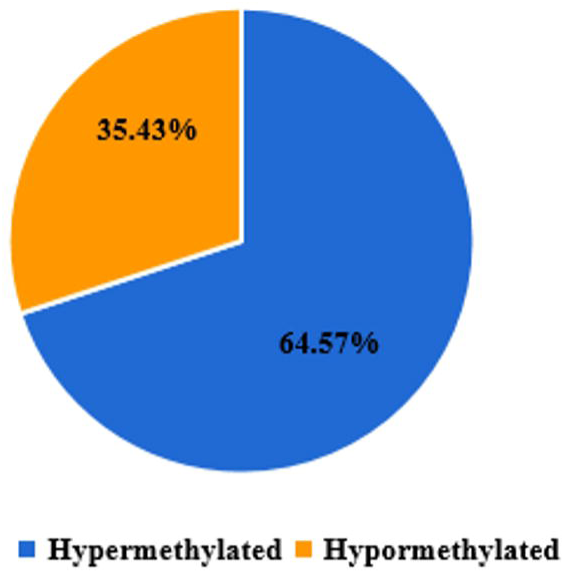
The proportion of hypermethylated and hypomethylated DMRs.

**Figure 2-Figure supplement 2.**
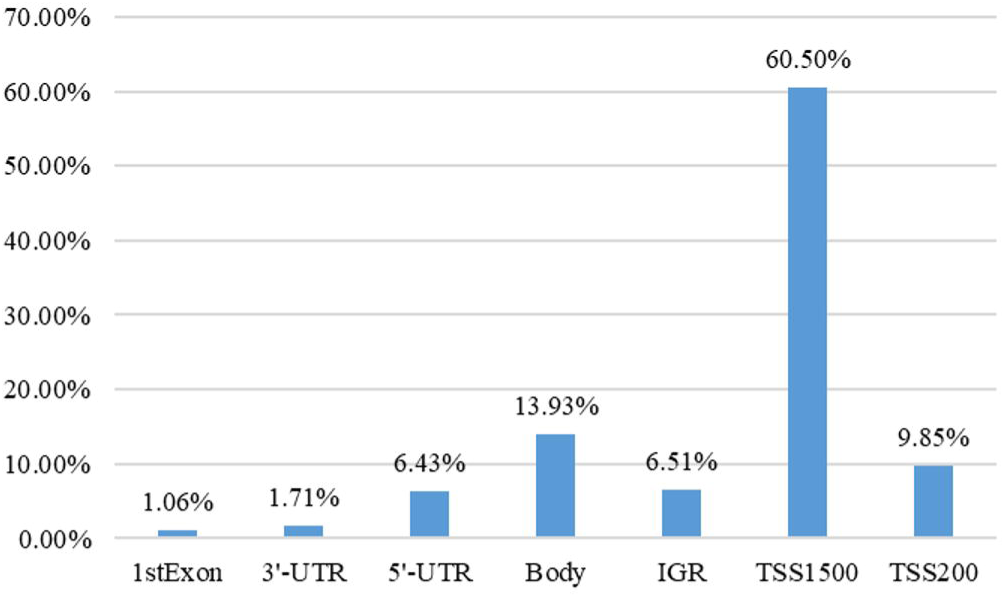
Proportion of DMRs for each genetic feature.

**Figure 2-Figure supplement 3.**
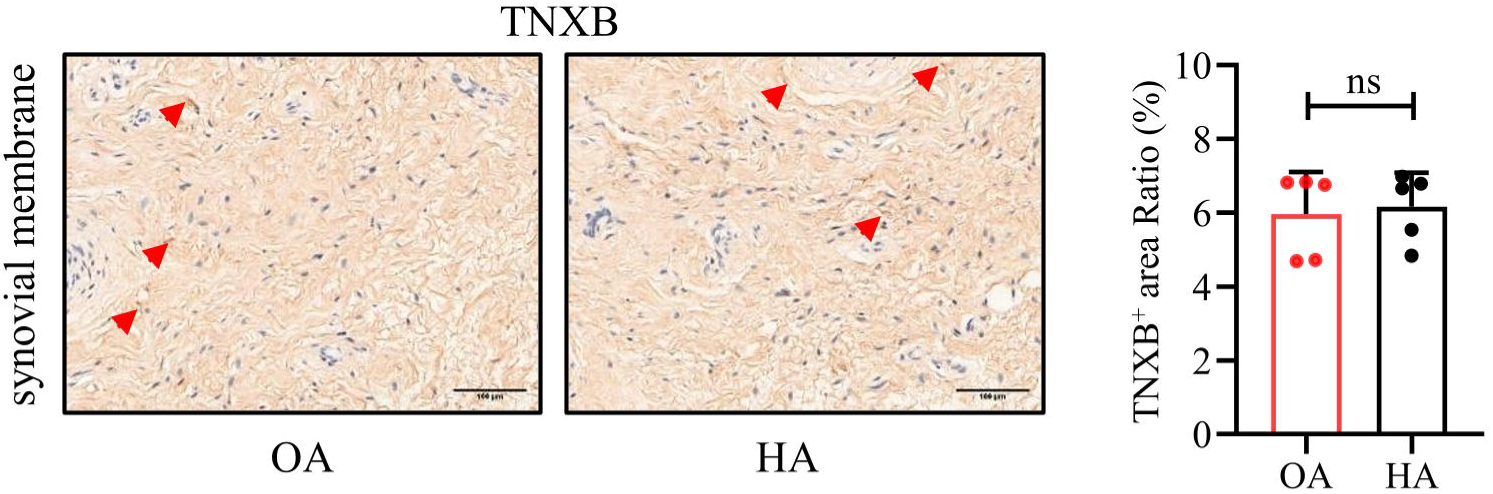
**(A)** Representative IHC staining of Tnxb in synovial membrane from human HA and osteoarthritis. Corresponding quantification of the proportion of **(B)** Tnxb positive regions. Red arrows indicate positive cells. Scale bar: 100 μm. Data were presented as means ± SD; n = 6 in each group. ns = no significance by unpaired Student’s t test.

**Figure 3-Figure supplement 1.**
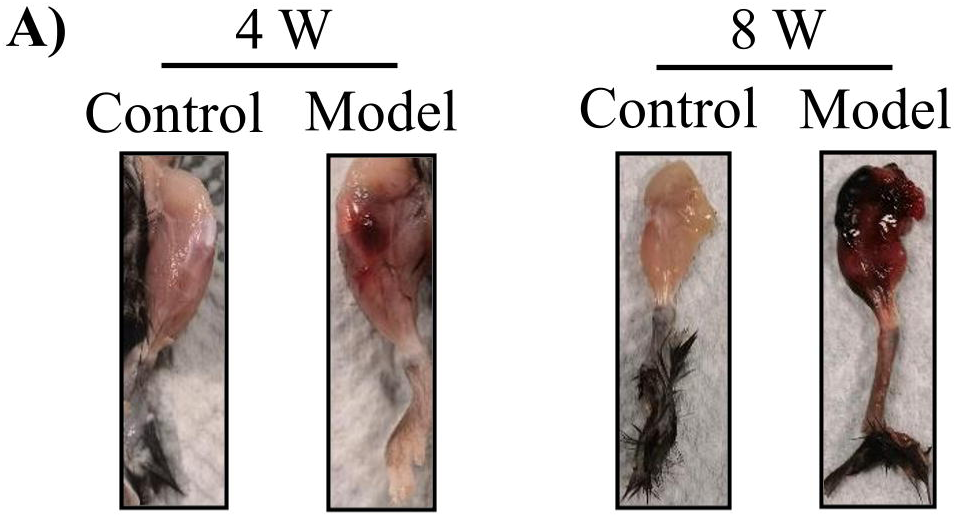
**(A)** Representative gross view of bleeding knee joint of *F8^-/-^* mice.

**Figure 3-Figure supplement 2.**
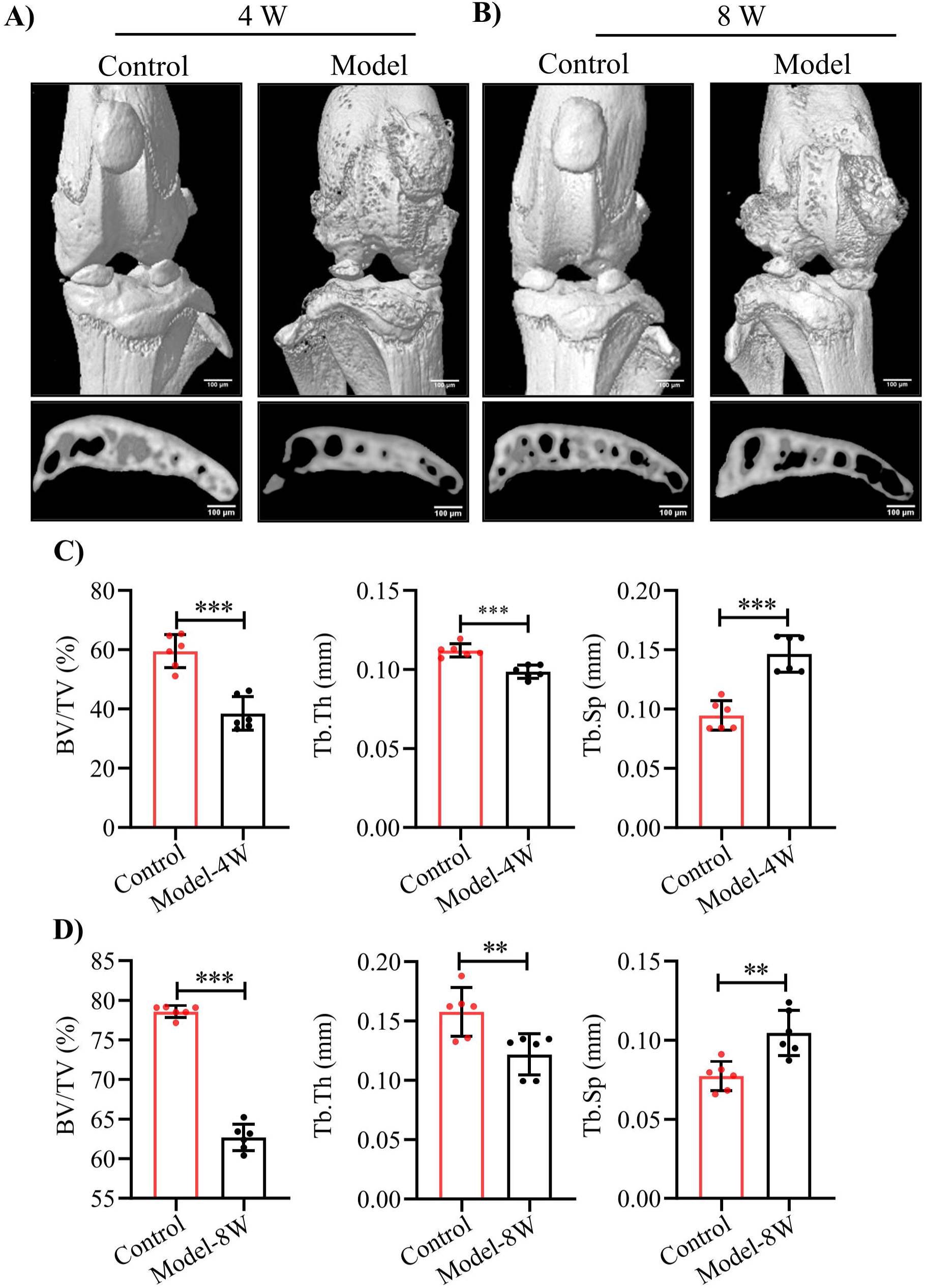
Representative 3D reconstruction of the knee joint and subchondral bone of HA model mice at **(A)** 4 weeks and **(B)** 8 weeks after puncture-induced injury. Quantification analysis of the subchondral bone BV/TV (%), Tb.Th (mm) and Tb.Sp (mm) at HA model mice at **(C)** 4 weeks and **(D)** 8 weeks after injury. 4 weeks. Data were presented as means ± SD and analyzed by 2-tailed unpaired parametric Student’s t test, \*\**P* < 0.01, \*\*\**P* < 0.001, n=6 in each group.

**Figure 4-Figure supplement 1.**
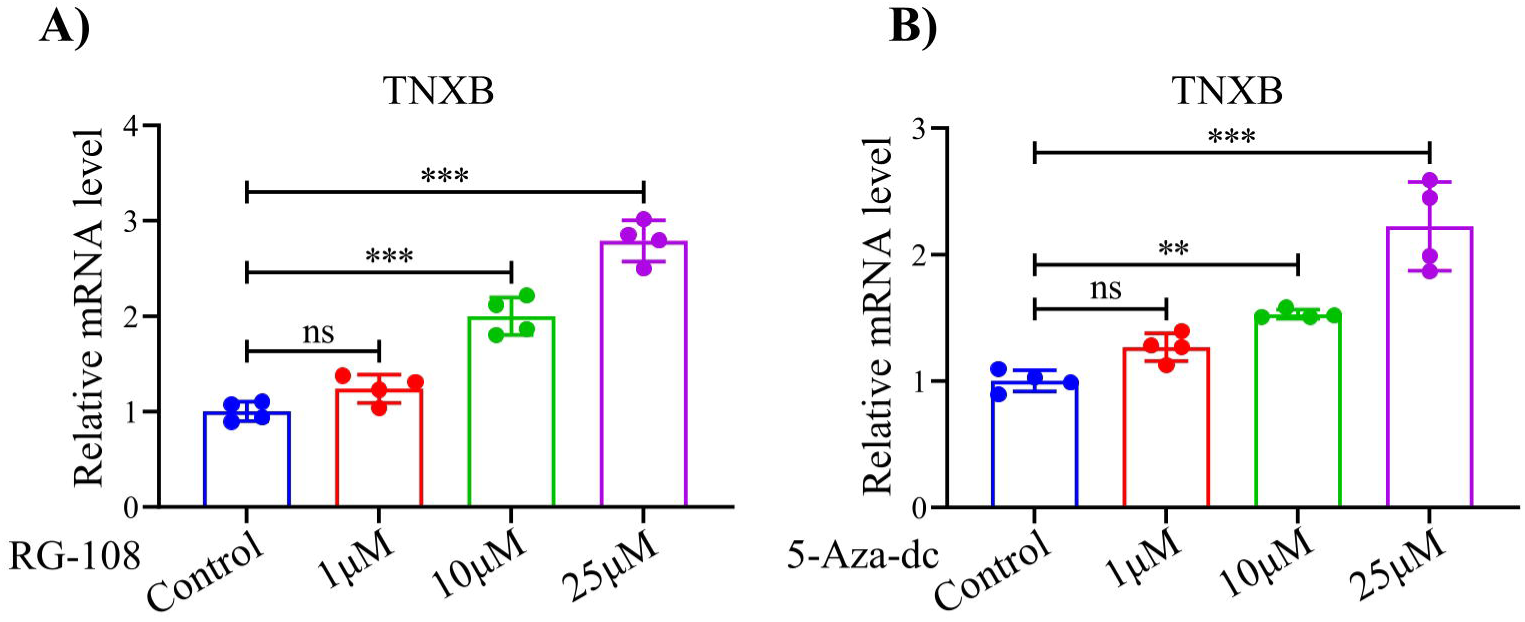
**(A)** and **(B)** qPCR analysis for Tnxb in mouse chondrocytes treated with RG-108 or 5-Aza-dc. Data were presented as means ± SD and analyzed by 2-tailed unpaired parametric Student’s t test, ns = no significance, \*\**P* < 0.01, \*\*\**P* < 0.001, n=4 in each group.

**Figure 4-Figure supplement 2.**
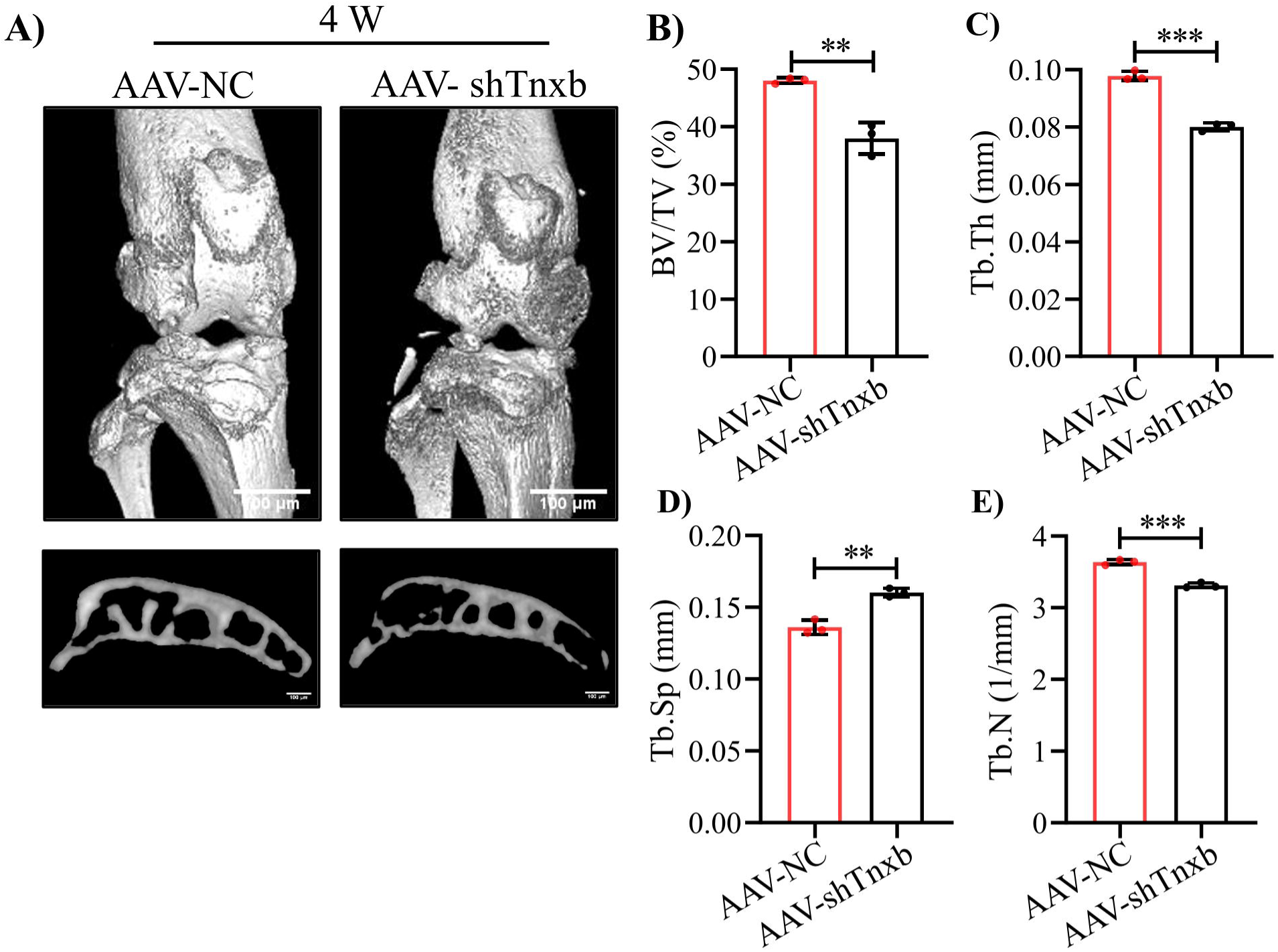
**(A)** Representative 3D reconstruction of the knee joint and subchondral bone of in Tnxb-KD HA mice. Quantification analysis of the subchondral bone **(B)** BV/TV (%), **(C)** Tb.Th (mm), **(D)** Tb.Sp (mm) and **(E)** Tb.N (1/mm). Data were presented as means ± SD and analyzed by 2-tailed unpaired parametric Student’s t test, \*\**P* < 0.01, \*\*\**P* < 0.001, n=3 in each group.

**Figure 5-Figure supplement 1.**
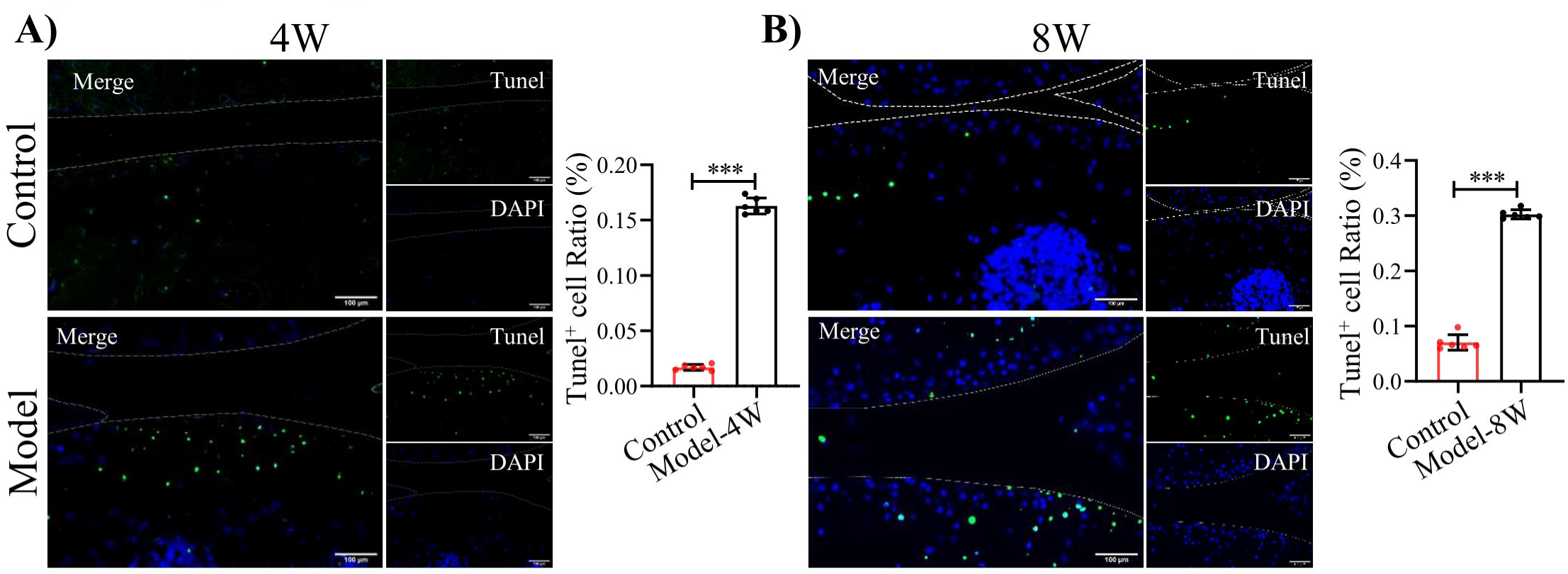
**(A)** TUNEL staining for apoptosis chondrocytes in articular cartilages from *F8^-/-^* mouse at 4 and 8 weeks after injury. Scale bar: 100 μm. **(B)** Quantification for TUNEL positive cells in articular cartilages.

**Figure 6-Figure supplement 1.**
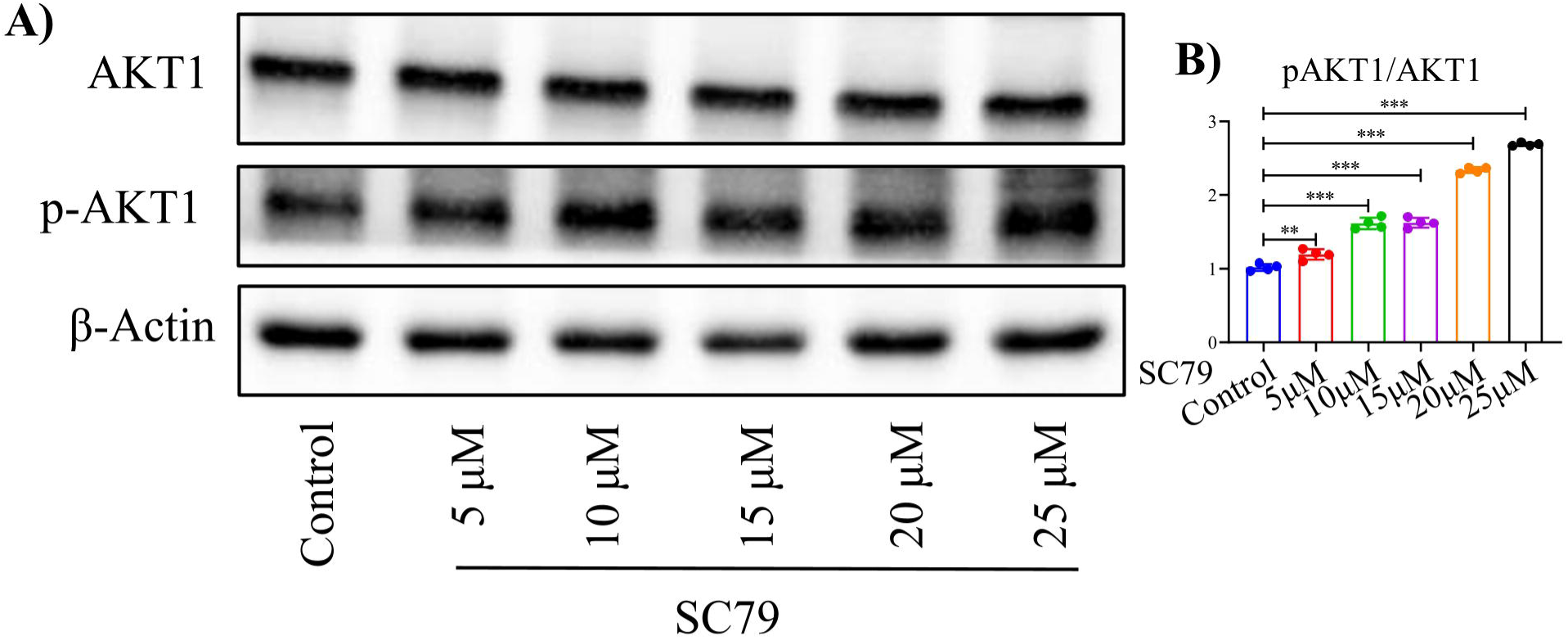
**(A)** Western blot examined the function of AKT agonist SC79. **(B)** Corresponding quantification of these proteins. Data were presented as means ± SD and analyzed by 2-tailed unpaired parametric Student’s t test, \*\**P* < 0.01, \*\*\**P* < 0.001, n = 4 in each group.

**Figure 6-Figure supplement 2.**
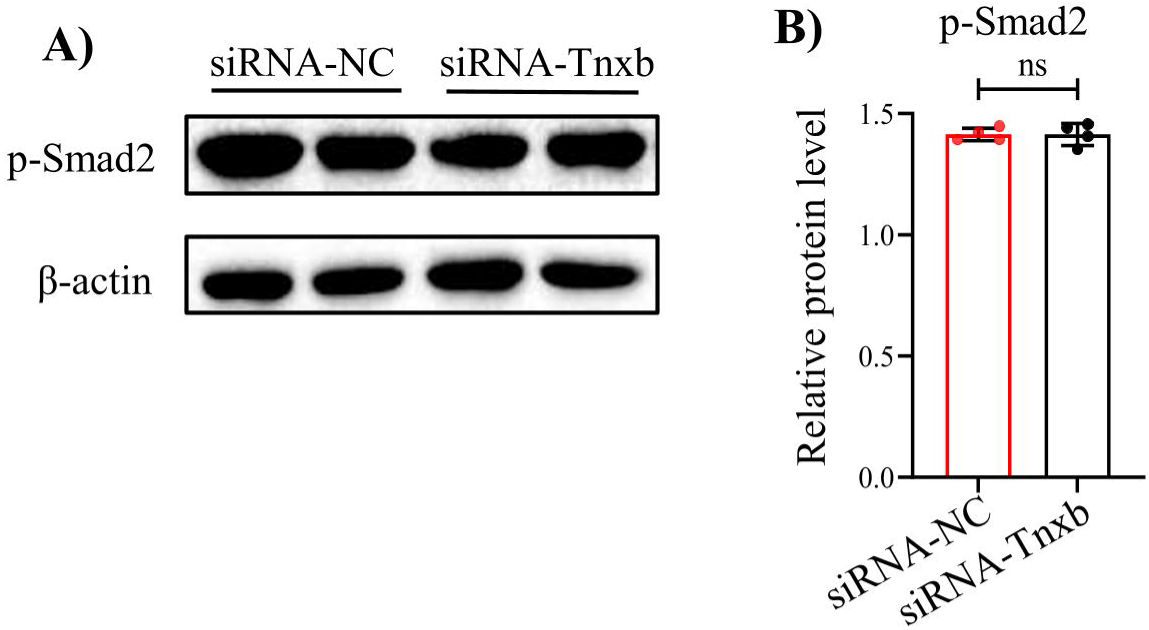
**(A)** Western blotting analysis for pSmad2 in mouse chondrocytes treated with siRNA-NC or siRNA-*Tnxb*. **(B)** Corresponding quantification of these proteins. Data were presented as means ± SD and analyzed by 2-tailed unpaired parametric Student’s t test, ns = no significance, n = 4 in each group.

**Supplement file 1.** Basic characteristics of subjects

**Supplement file 2.** Primer sequences for qPCR

**Supplement file 3a**. DMRs between HA and OA cartilages

**Supplement file 3b.** GO pathway enrichment analysis of DMRs-related gene (Top 10 pathways)

**Supplement file 3c.** KEGG pathway enrichment analysis of DMRs-related gene (Top 10 pathways)

**Figure 4-source data 1**

Original file for the Western blot analysis of β-actin in Figure 4B

**Figure 4-source data 2**

Original file for the Western blot analysis of Tnxb in Figure 4B

**Figure 4-source data 3**

PDF containing Figure 4B and original scans of the relevant Western blot analysis (anti-β-actin, and anti-Tnxb) with highlighted bands and sample labels

**Figure 4-source data 4**

Original file for the Western blot analysis of β-actin in Figure 4E

**Figure 4-source data 5**

Original file for the Western blot analysis of Col2a1 in Figure 4E

**Figure 4-source data 6**

Original file for the Western blot analysis of Mmp13 in Figure 4E

**Figure 4-source data 7**

PDF containing Figure 4E and original scans of the relevant Western blot analysis (anti-β-actin, anti-Col2a1, and anti-Mmp13) with highlighted bands and sample labels

**Figure 5-source data 1**

Original file for the Western blot analysis of β-actin in Figure 5C

**Figure 5-source data 2**

Original file for the Western blot analysis of Bax in Figure 5C

**Figure 5-source data 3**

Original file for the Western blot analysis of Bcl-2 in Figure 5C

**Figure 5-source data 4**

Original file for the Western blot analysis of cleaved-caspase 3 in Figure 5C

**Figure 5-source data 5**

PDF containing Figure 5C and original scans of the relevant Western blot analysis (anti-β-actin, anti-Bax, anti-Bcl-2, and anti-cleaved-caspase 3) with highlighted bands and sample labels

**Figure 6—source data 1**

Original file for the Western blot analysis of β-actin in Figure 6A

**Figure 6—source data 2**

Original file for the Western blot analysis of Col2a1 in Figure 6A

**Figure 6—source data 3**

Original file for the Western blot analysis of Mmp13 in Figure 6A

**Figure 6—source data 4**

Original file for the Western blot analysis of AKT1 in Figure 6A

**Figure 6—source data 5**

Original file for the Western blot analysis of pAKT1 in Figure 6A

**Figure 6—source data 6**

Original file for the Western blot analysis of Bax in Figure 6A

**Figure 6—source data 7**

Original file for the Western blot analysis of Bcl-2 in Figure 6A

**Figure 6—source data 8**

Original file for the Western blot analysis of cleaved-caspase9 in Figure 6A

**Figure 6—source data 9**

Original file for the Western blot analysis of cleaved-caspase9-β-actin in Figure 6A

**Figure 6—source data 10**

PDF containing Figure 6A and original scans of the relevant Western blot analysis (anti-β-actin, anti-Col2a1, anti-Mmp13, anti-AKT1, anti-pAKT1, anti-Bax, anti-Bcl-2, anti-cleaved-caspase 9, and anti-cleaved-caspase 9-β-actin) with highlighted bands and sample labels

**Figure 6-Figure supplement 1—source data 1**

Original file for the Western blot analysis of β-actin in Figure 6-Figure supplement 1

**Figure 6-Figure supplement 1—source data 2**

Original file for the Western blot analysis of AKT1 in Figure 6-Figure supplement 1

**Figure 6-Figure supplement 1—source data 3**

Original file for the Western blot analysis of pAKT1 in Figure 6-Figure supplement 1

**Figure 6-Figure supplement 1—source data 4**

PDF containing Figure 6-Figure supplement 1 and original scans of the relevant Western blot analysis (anti-β-actin, anti-AKT1, and anti-pAKT1) with highlighted bands and sample labels

**Figure 6-Figure supplement 2—source data 1**

Original file for the Western blot analysis of β-actin in Figure 6-Figure supplement 2

**Figure 6-Figure supplement 2—source data 2**

Original file for the Western blot analysis of pSmad2 in Figure 6-Figure supplement 2

**Figure 6-Figure supplement 1—source data 2**

PDF containing Figure 6-Figure supplement 2 and original scans of the relevant Western blot analysis (anti-β-actin, and anti-pSmad2) with highlighted bands and sample labels

## References

1. Carulli, C., Innocenti, M., Linari, S., Morfini, M., Castaman, G., and Innocenti, M. (2021). Joint replacement for the management of haemophilic arthropathy in patients with inhibitors: A long-term experience at a single Haemophilia centre. Haemophilia 27, e93–e101. 10.1111/hae.14169.

2. Gualtierotti, R., Solimeno, L.P., and Peyvandi, F. (2021). Hemophilic arthropathy: Current knowledge and future perspectives. J Thromb Haemost 19, 2112–2121. 10.1111/jth.15444.

3. Di Minno, M.N., Ambrosino, P., Franchini, M., Coppola, A., and Di Minno, G. (2013). Arthropathy in patients with moderate hemophilia a: a systematic review of the literature. Semin Thromb Hemost 39, 723–731. 10.1055/s-0033-1354422.

4. Wyseure, T., Yang, T., Zhou, J.Y., Cooke, E.J., Wanko, B., Olmer, M., Agashe, R., Morodomi, Y., Behrendt, N., Lotz, M., et al. (2019). TAFI deficiency causes maladaptive vascular remodeling after hemophilic joint bleeding. JCI Insight 4. 10.1172/jci.insight.128379.

5. Aledort, L.M., Haschmeyer, R.H., and Pettersson, H. (1994). A longitudinal study of orthopaedic outcomes for severe factor-VIII-deficient haemophiliacs. The Orthopaedic Outcome Study Group. J Intern Med 236, 391–399. 10.1111/j.1365-2796.1994.tb00815.x.

6. Bolton-Maggs, P.H., and Pasi, K.J. (2003). Haemophilias A and B. Lancet 361, 1801–1809. 10.1016/S0140-6736(03)13405-8.

7. Fischer, K., Bom, J.G., Mauser-Bunschoten, E.P., Roosendaal, G., and Berg, H.M. (2005). Effects of haemophilic arthropathy on health-related quality of life and socio-economic parameters. Haemophilia 11, 43–48. 10.1111/j.1365-2516.2005.01065.x.

8. Manco-Johnson, M.J., Abshire, T.C., Shapiro, A.D., Riske, B., Hacker, M.R., Kilcoyne, R., Ingram, J.D., Manco-Johnson, M.L., Funk, S., Jacobson, L., et al. (2007). Prophylaxis versus episodic treatment to prevent joint disease in boys with severe hemophilia. N Engl J Med 357, 535–544. 10.1056/NEJMoa067659.

9. Oldenburg, J. (2015). Optimal treatment strategies for hemophilia: achievements and limitations of current prophylactic regimens. Blood 125, 2038–2044. 10.1182/blood-2015-01-528414.

10. Olivieri, M., Kurnik, K., Pfluger, T., and Bidlingmaier, C. (2012). Identification and long-term observation of early joint damage by magnetic resonance imaging in clinically asymptomatic joints in patients with haemophilia A or B despite prophylaxis. Haemophilia 18, 369–374. 10.1111/j.1365-2516.2011.02682.x.

11. Jackson, S.C., Yang, M., Minuk, L., Sholzberg, M., St-Louis, J., Iorio, A., Card, R., and Poon, M.C. (2015). Prophylaxis in older Canadian adults with hemophilia A: lessons and more questions. BMC Hematol 15, 4. 10.1186/s12878-015-0022-8.

12. Schramm, W., Gringeri, A., Ljung, R., Berger, K., Crispin, A., Bullinger, M., Giangrande, P.L., Von Mackensen, S., Mantovani, L.G., Nemes, L., et al. (2012). Haemophilia care in Europe: the ESCHQoL study. Haemophilia 18, 729–737. 10.1111/j.1365-2516.2012.02847.x.

13. Kleiboer, B., Layer, M.A., Cafuir, L.A., Cuker, A., Escobar, M., Eyster, M.E., Kraut, E., Leavitt, A.D., Lentz, S.R., Quon, D., et al. (2022). Postoperative bleeding complications in patients with hemophilia undergoing major orthopedic surgery: A prospective multicenter observational study. J Thromb Haemost 20, 857–865. 10.1111/jth.15654.

14. Wojdasiewicz, P., Poniatowski, L.A., Nauman, P., Mandat, T., Paradowska-Gorycka, A., Romanowska-Prochnicka, K., Szukiewicz, D., Kotela, A., Kubaszewski, L., Kotela, I., et al. (2018). Cytokines in the pathogenesis of hemophilic arthropathy. Cytokine Growth Factor Rev 39, 71–91. 10.1016/j.cytogfr.2017.11.003.

15. Pulles, A.E., Mastbergen, S.C., Schutgens, R.E., Lafeber, F.P., and van Vulpen, L.F. (2017). Pathophysiology of hemophilic arthropathy and potential targets for therapy. Pharmacol Res 115, 192–199. 10.1016/j.phrs.2016.11.032.

16. Fermor, B., Christensen, S.E., Youn, I., Cernanec, J.M., Davies, C.M., and Weinberg, J.B. (2007). Oxygen, nitric oxide and articular cartilage. Eur Cell Mater 13, 56–65; discussion 65. 10.22203/ecm.v013a06.

17. Lafont, J.E. (2010). Lack of oxygen in articular cartilage: consequences for chondrocyte biology. Int J Exp Pathol 91, 99–106. 10.1111/j.1365-2613.2010.00707.x.

18. van Vulpen, L.F.D., Holstein, K., and Martinoli, C. (2018). Joint disease in haemophilia: Pathophysiology, pain and imaging. Haemophilia 24 *Suppl 6*, 44–49. 10.1111/hae.13449.

19. Feil, R., and Fraga, M.F. (2012). Epigenetics and the environment: emerging patterns and implications. Nat Rev Genet 13, 97–109. 10.1038/nrg3142.

20. Jirtle, R.L., and Skinner, M.K. (2007). Environmental epigenomics and disease susceptibility. Nat Rev Genet 8, 253–262. 10.1038/nrg2045.

21. Stelzer, Y., Shivalila, C.S., Soldner, F., Markoulaki, S., and Jaenisch, R. (2015). Tracing dynamic changes of DNA methylation at single-cell resolution. Cell 163, 218–229. 10.1016/j.cell.2015.08.046.

22. Wang, W., Yu, Y., Hao, J., Wen, Y., Han, J., Hou, W., Liu, R., Zhao, B., He, A., Li, P., et al. (2017). Genome-wide DNA methylation profiling of articular cartilage reveals significant epigenetic alterations in Kashin-Beck disease and osteoarthritis. Osteoarthritis Cartilage 25, 2127–2133. 10.1016/j.joca.2017.08.002.

23. Zhang, Y., Fukui, N., Yahata, M., Katsuragawa, Y., Tashiro, T., Ikegawa, S., and Lee, M.T. (2016). Genome-wide DNA methylation profile implicates potential cartilage regeneration at the late stage of knee osteoarthritis. Osteoarthritis Cartilage 24, 835–843. 10.1016/j.joca.2015.12.013.

24. Cooke, E.J., Zhou, J.Y., Wyseure, T., Joshi, S., Bhat, V., Durden, D.L., Mosnier, L.O., and von Drygalski, A. (2018). Vascular Permeability and Remodelling Coincide with Inflammatory and Reparative Processes after Joint Bleeding in Factor VIII-Deficient Mice. Thromb Haemost 118, 1036–1047. 10.1055/s-0038-1641755.

25. Kalebota, N., Salai, G., Peric, P., Hrkac, S., Novak, R., Durmis, K.K., and Grgurevic, L. (2022). ADAMTS-4 as a possible distinguishing indicator between osteoarthritis and haemophilic arthropathy. Haemophilia 28, 656–662. 10.1111/hae.14569.

26. Hakobyan, N., Enockson, C., Cole, A.A., Sumner, D.R., and Valentino, L.A. (2008). Experimental haemophilic arthropathy in a mouse model of a massive haemarthrosis: gross, radiological and histological changes. Haemophilia 14, 804–809. 10.1111/j.1365-2516.2008.01689.x.

27. Chen, Y.R., Yang, K.C., Lu, D.H., Wu, W.T., Wang, C.C., and Tsai, M.H. (2019). The chondroprotective effect of diosmin on human articular chondrocytes under oxidative stress. Phytother Res 33, 2378–2386. 10.1002/ptr.6425.

28. Maly, K., Andres Sastre, E., Farrell, E., Meurer, A., and Zaucke, F. (2021). COMP and TSP-4: Functional Roles in Articular Cartilage and Relevance in Osteoarthritis. Int J Mol Sci 22. 10.3390/ijms22052242.

29. Zhang, H., Chen, C., Cui, Y., Li, Y., Wang, Z., Mao, X., Dou, P., Li, Y., and Ma, C. (2019). lnc-SAMD14-4 can regulate expression of the COL1A1 and COL1A2 in human chondrocytes. PeerJ 7, e7491. 10.7717/peerj.7491.

30. Haxaire, C., Hakobyan, N., Pannellini, T., Carballo, C., McIlwain, D., Mak, T.W., Rodeo, S., Acharya, S., Li, D., Szymonifka, J., et al. (2018). Blood-induced bone loss in murine hemophilic arthropathy is prevented by blocking the iRhom2/ADAM17/TNF-alpha pathway. Blood 132, 1064–1074. 10.1182/blood-2017-12-820571.

31. Magisetty, J., Kondreddy, V., Keshava, S., Das, K., Esmon, C.T., Pendurthi, U.R., and Rao, L.V.M. (2022). Selective inhibition of activated protein C anticoagulant activity protects against hemophilic arthropathy in mice. Blood 139, 2830–2841. 10.1182/blood.2021013119.

32. He, Y., Zheng, Z., Liu, C., Li, W., Zhao, L., Nie, G., and Li, H. (2022). Inhibiting DNA methylation alleviates cisplatin-induced hearing loss by decreasing oxidative stress-induced mitochondria-dependent apoptosis via the LRP1-PI3K/AKT pathway. Acta Pharm Sin B 12, 1305–1321. 10.1016/j.apsb.2021.11.002.

33. Zhu, X., Chen, F., Lu, K., Wei, A., Jiang, Q., and Cao, W. (2019). PPARgamma preservation via promoter demethylation alleviates osteoarthritis in mice. Ann Rheum Dis 78, 1420–1429. 10.1136/annrheumdis-2018-214940.

34. Cai, C., Min, S., Yan, B., Liu, W., Yang, X., Li, L., Wang, T., and Jin, A. (2019). MiR-27a promotes the autophagy and apoptosis of IL-1beta treated-articular chondrocytes in osteoarthritis through PI3K/AKT/mTOR signaling. Aging (Albany NY) 11, 6371–6384. 10.18632/aging.102194.

35. Hu, P.F., Chen, W.P., Bao, J.P., and Wu, L.D. (2018). Paeoniflorin inhibits IL-1beta-induced chondrocyte apoptosis by regulating the Bax/Bcl-2/caspase-3 signaling pathway. Mol Med Rep 17, 6194–6200. 10.3892/mmr.2018.8631.

36. Luo, J., Manning, B.D., and Cantley, L.C. (2003). Targeting the PI3K-Akt pathway in human cancer: rationale and promise. Cancer Cell 4, 257–262. 10.1016/s1535-6108(03)00248-4.

37. Hooiveld, M.J., Roosendaal, G., van den Berg, H.M., Bijlsma, J.W., and Lafeber, F.P. (2003). Haemoglobin-derived iron-dependent hydroxyl radical formation in blood-induced joint damage: an in vitro study. Rheumatology (Oxford) 42, 784–790. 10.1093/rheumatology/keg220.

38. Srivastava, A. (2015). Inflammation is key to hemophilic arthropathy. Blood 126, 2175–2176. 10.1182/blood-2015-09-665091.

39. van Vulpen, L.F., van Meegeren, M.E., Roosendaal, G., Jansen, N.W., van Laar, J.M., Schutgens, R.E., Mastbergen, S.C., and Lafeber, F.P. (2015). Biochemical markers of joint tissue damage increase shortly after a joint bleed; an explorative human and canine in vivo study. Osteoarthritis Cartilage 23, 63–69. 10.1016/j.joca.2014.09.008.

40. Visser, A.W., de Mutsert, R., le Cessie, S., den Heijer, M., Rosendaal, F.R., Kloppenburg, M., and Group, N.E.O.S. (2015). The relative contribution of mechanical stress and systemic processes in different types of osteoarthritis: the NEO study. Ann Rheum Dis 74, 1842–1847. 10.1136/annrheumdis-2013-205012.

41. Iijima, H., Gilmer, G., Wang, K., Bean, A.C., He, Y., Lin, H., Tang, W.Y., Lamont, D., Tai, C., Ito, A., et al. (2023). Age-related matrix stiffening epigenetically regulates alpha-Klotho expression and compromises chondrocyte integrity. Nat Commun 14, 18. 10.1038/s41467-022-35359-2.

42. Kucuk, C., Hu, X., Jiang, B., Klinkebiel, D., Geng, H., Gong, Q., Bouska, A., Iqbal, J., Gaulard, P., McKeithan, T.W., and Chan, W.C. (2015). Global promoter methylation analysis reveals novel candidate tumor suppressor genes in natural killer cell lymphoma. Clin Cancer Res 21, 1699–1711. 10.1158/1078-0432.CCR-14-1216.

43. Kulis, M., and Esteller, M. (2010). DNA methylation and cancer. Adv Genet 70, 27–56. 10.1016/B978-0-12-380866-0.60002-2.

44. Jones, F.S., and Jones, P.L. (2000). The tenascin family of ECM glycoproteins: structure, function, and regulation during embryonic development and tissue remodeling. Dev Dyn 218, 235–259. 10.1002/(SICI)1097-0177(200006)218:2<235::AID-DVDY2>3.0.CO;2-G.

45. Kim, S.H., Turnbull, J., and Guimond, S. (2011). Extracellular matrix and cell signalling: the dynamic cooperation of integrin, proteoglycan and growth factor receptor. J Endocrinol 209, 139–151. 10.1530/JOE-10-0377.

46. Alcaraz, L.B., Exposito, J.Y., Chuvin, N., Pommier, R.M., Cluzel, C., Martel, S., Sentis, S., Bartholin, L., Lethias, C., and Valcourt, U. (2014). Tenascin-X promotes epithelial-to-mesenchymal transition by activating latent TGF-beta. J Cell Biol 205, 409–428. 10.1083/jcb.201308031.

47. Sutherland, A.J., Converse, G.L., Hopkins, R.A., and Detamore, M.S. (2015). The bioactivity of cartilage extracellular matrix in articular cartilage regeneration. Adv Healthc Mater 4, 29–39. 10.1002/adhm.201400165.

48. Anaparti, V., Agarwal, P., Smolik, I., Mookherjee, N., and El-Gabalawy, H. (2020). Whole Blood Targeted Bisulfite Sequencing and Differential Methylation in the C6ORF10 Gene of Patients with Rheumatoid Arthritis. J Rheumatol 47, 1614–1623. 10.3899/jrheum.190376.

49. Porter, L.F., Saptarshi, N., Fang, Y., Rathi, S., den Hollander, A.I., de Jong, E.K., Clark, S.J., Bishop, P.N., Olsen, T.W., Liloglou, T., et al. (2019). Whole-genome methylation profiling of the retinal pigment epithelium of individuals with age-related macular degeneration reveals differential methylation of the SKI, GTF2H4, and TNXB genes. Clin Epigenetics 11, 6. 10.1186/s13148-019-0608-2.

50. Tucker, R.P., Drabikowski, K., Hess, J.F., Ferralli, J., Chiquet-Ehrismann, R., and Adams, J.C. (2006). Phylogenetic analysis of the tenascin gene family: evidence of origin early in the chordate lineage. BMC Evol Biol 6, 60. 10.1186/1471-2148-6-60.

51. Hasegawa, M., Yoshida, T., and Sudo, A. (2018). Role of tenascin-C in articular cartilage. Mod Rheumatol 28, 215–220. 10.1080/14397595.2017.1349560.

52. Hasegawa, M., Yoshida, T., and Sudo, A. (2020). Tenascin-C in Osteoarthritis and Rheumatoid Arthritis. Front Immunol 11, 577015. 10.3389/fimmu.2020.577015.

53. Valcourt, U., Alcaraz, L.B., Exposito, J.Y., Lethias, C., and Bartholin, L. (2015). Tenascin-X: beyond the architectural function. Cell Adh Migr 9, 154–165. 10.4161/19336918.2014.994893.

54. Hooiveld, M., Roosendaal, G., Wenting, M., van den Berg, M., Bijlsma, J., and Lafeber, F. (2003). Short-term exposure of cartilage to blood results in chondrocyte apoptosis. Am J Pathol 162, 943–951. 10.1016/S0002-9440(10)63889-8.

55. Jang, J.H., and Chung, C.P. (2005). Tenascin-C promotes cell survival by activation of Akt in human chondrosarcoma cell. Cancer Lett 229, 101–105. 10.1016/j.canlet.2004.12.012.

56. Tan, B.S., Tiong, K.H., Choo, H.L., Chung, F.F., Hii, L.W., Tan, S.H., Yap, I.K., Pani, S., Khor, N.T., Wong, S.F., et al. (2015). Mutant p53-R273H mediates cancer cell survival and anoikis resistance through AKT-dependent suppression of BCL2-modifying factor (BMF). Cell Death Dis 6, e1826. 10.1038/cddis.2015.191.

57. Testa, J.R., and Bellacosa, A. (2001). AKT plays a central role in tumorigenesis. Proc Natl Acad Sci U S A 98, 10983–10985. 10.1073/pnas.211430998.

58. Zheng, L., Yao, Y., Luo, D., Han, Z., Zhang, X., Pang, N., Ding, M., Ye, H., Zhu, K., and Yi, W. (2023). Iron regulates chondrocyte phenotype in haemophilic cartilage through the PTEN/PI3 K/AKT/FOXO1 pathway. Hematology 28, 2240585. 10.1080/16078454.2023.2240585.

59. El Mabrouk, M., Sylvester, J., and Zafarullah, M. (2007). Signaling pathways implicated in oncostatin M-induced aggrecanase-1 and matrix metalloproteinase-13 expression in human articular chondrocytes. Biochim Biophys Acta 1773, 309–320. 10.1016/j.bbamcr.2006.11.018.

60. Lu, J., Feng, X., Zhang, H., Wei, Y., Yang, Y., Tian, Y., and Bai, L. (2021). Maresin-1 suppresses IL-1beta-induced MMP-13 secretion by activating the PI3K/AKT pathway and inhibiting the NF-kappaB pathway in synovioblasts of an osteoarthritis rat model with treadmill exercise. Connect Tissue Res 62, 508–518. 10.1080/03008207.2020.1780218.

