## Supplement file 1 for "Aberrant methylation and expression of TNXB promote chondrocyte apoptosis and extracullar matrix degradation in hemophilic arthropathy via AKT signaling"

**Supplement file 1.** Basic characteristics of subjects

| **Sample ID** | **Age** | **Gender** | **Height (cm)** | **Weight (kg)** | **BMI** |
| --- | --- | --- | --- | --- | --- |
| HA1 | 43 | male | 165.20 | 65.80 | 24.17 |
| HA2 | 43 | male | 178.30 | 95.10 | 30.02 |
| HA3 | 50 | male | 159.10 | 58.20 | 23.02 |
| HA4 | 40 | male | 162.50 | 58.30 | 22.21 |
| HA5 | 37 | male | 168.50 | 60.50 | 21.44 |
| OA1 | 66 | male | 166.40 | 64.20 | 23.30 |
| OA2 | 60 | male | 162.60 | 71.00 | 27.05 |
| OA3 | 68 | female | 162.40 | 61.10 | 23.28 |
| OA4 | 70 | female | 155.30 | 70.20 | 29.22 |
| OA5 | 71 | female | 149.80 | 62.60 | 28.20 |
