## Supplement file 2 for "Aberrant methylation and expression of TNXB promote chondrocyte apoptosis and extracullar matrix degradation in hemophilic arthropathy via AKT signaling"

**Supplement file 2.** Primer sequences for qPCR

| **Primer name** | **Primer sequence (5’→3’)** |
| --- | --- |
| *Col2a1* Forward | GGGAATGTCCTCTGCGATGAC |
| *Col2a1* Reverse | GAAGGGGATCTCGGGGTTG |
| *Mmp13* Forward | CTTCTTCTTGTTGAGCTGGACTC |
| *Mmp13* Reverse | CTGTGGAGGTCACTGTAGACT |
| *Tnxb* Forward | TCCGTGTAGACTCAGCAAAGG |
| *Tnxb* Reverse | CCCCACGATAAGAGACAGCG |
| *Actb* Forward | GGCTGTATTCCCCTCCATCG |
| *Actb* Reverse | CCAGTTGGTAACAATGCCATGT |
