## Supplementary figures and images for "Aberrant methylation and expression of TNXB promote chondrocyte apoptosis and extracullar matrix degradation in hemophilic arthropathy via AKT signaling"

### Figure 4—source data 3.pdf

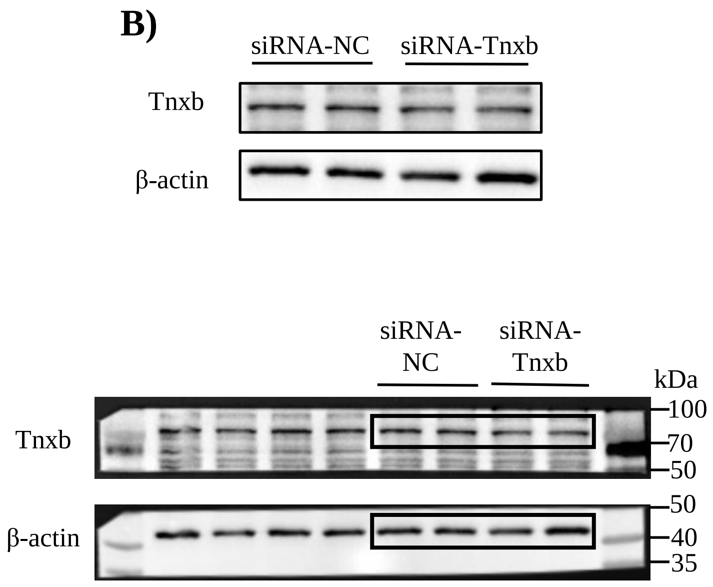

### Figure 4—source data 7.pdf

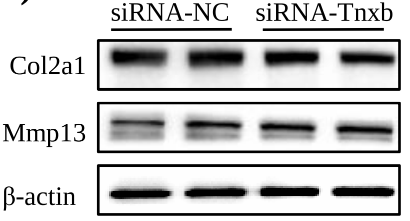

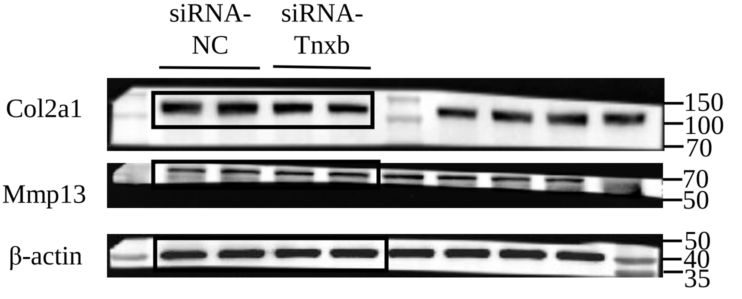

### Figure 5—source data 5.pdf

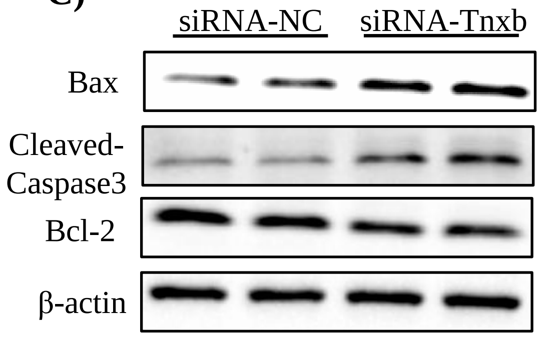

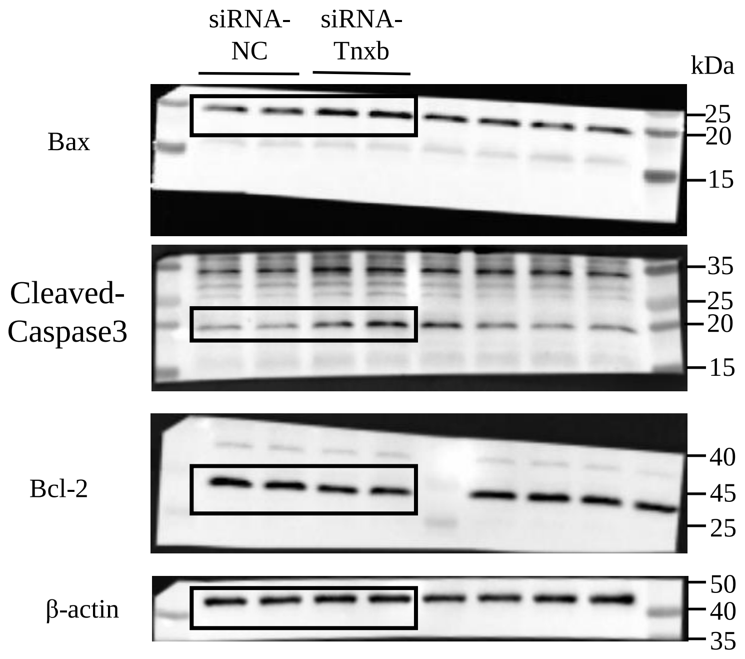

### Figure 6-Figure supplement 1—source data 4.pdf

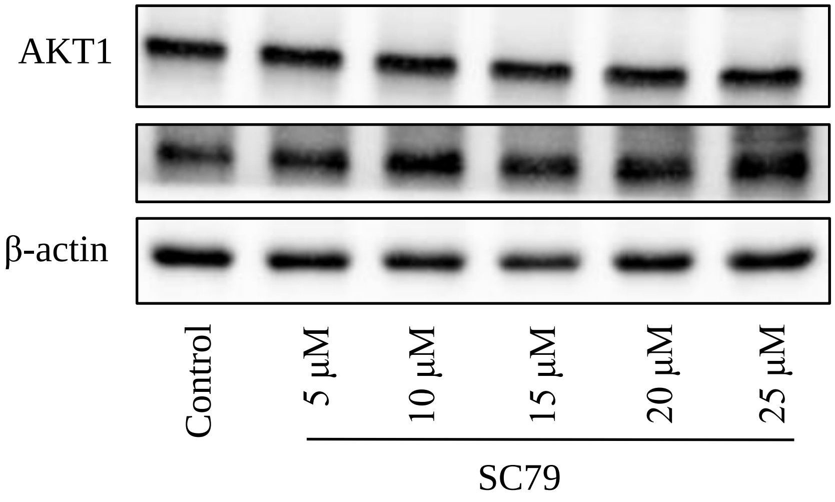

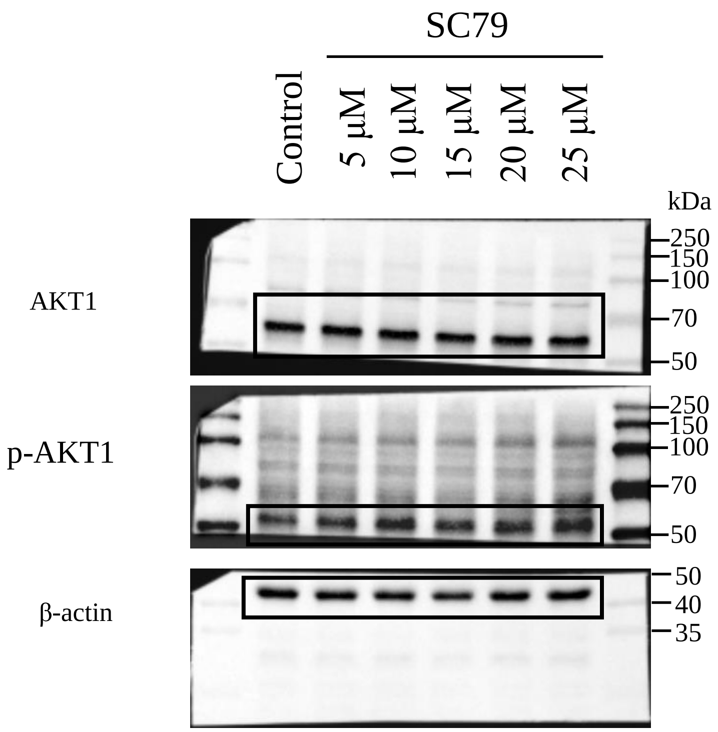

### Figure 6-Figure supplement 2—source data 3.pdf

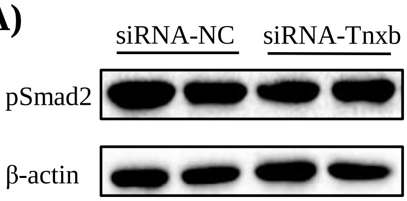

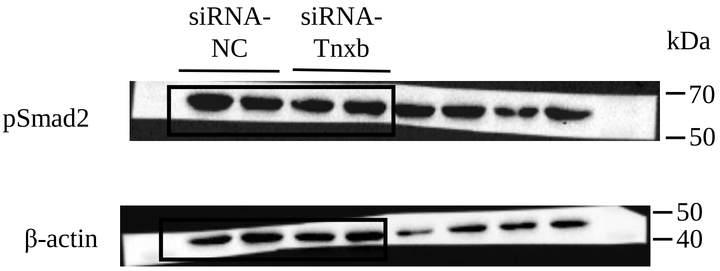

### Figure 6—source data 10.pdf

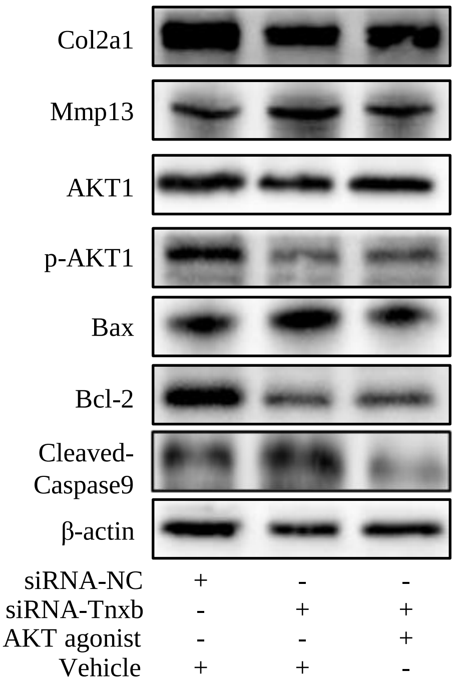

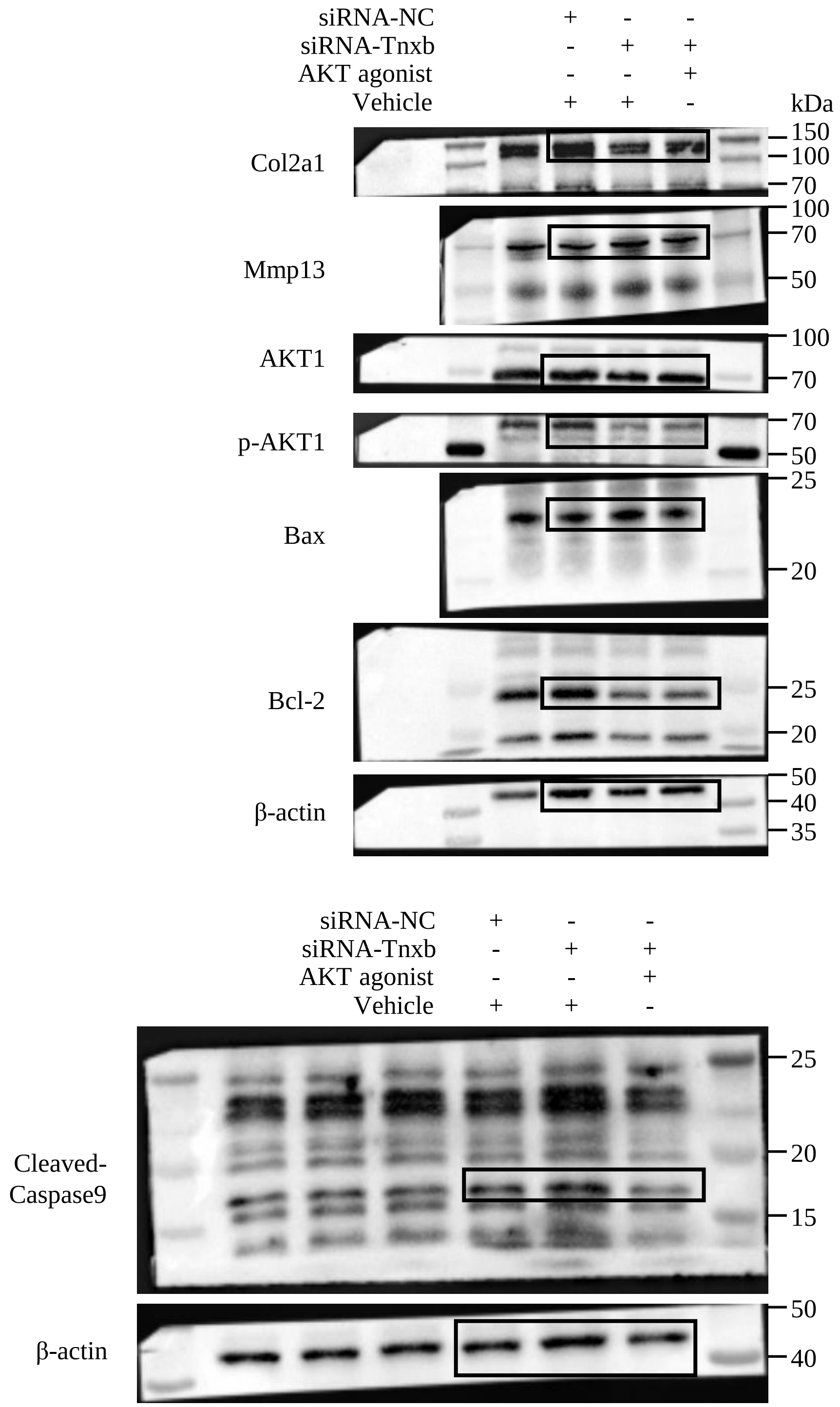
